# Sequence-based GWAS in 180 000 German Holstein cattle reveals new candidate genes for milk production traits

**DOI:** 10.1101/2023.12.06.570350

**Authors:** Ana-Marija Križanac, Christian Reimer, Johannes Heise, Zengting Liu, Jennie Pryce, Jörn Bennewitz, Georg Thaller, Clemens Falker-Gieske, Jens Tetens

## Abstract

**Background:** The use of genome-wide association studies (GWAS) has led to the identification of numerous quantitative trait loci and candidate genes in dairy cattle. To obtain sufficient power of GWAS and to identify quantitative trait nucleotides, whole-genome sequence data is required. Sequence data facilitates the identification of potential causal variants; however, sequencing of whole genomes is still expensive for a large number of animals. Imputation is a quick and efficient way of obtaining sequence data from large datasets. Milk production traits are complex and influenced by many genetic and environmental factors. Although extensive research has been performed for these traits, with many associations unveiled thus far, due to their crucial economic importance, complex genetic architecture, and the fact that causative variants in cattle are still scarce, there is a need for a better understanding of their genetic background. In this study, we aimed to identify new candidate loci associated with milk production traits in German Holstein cattle, the most important dairy breed in Germany and worldwide. For that purpose, 252,285 cattle were imputed to the sequence level and large-scale GWAS was carried out to identify new association signals.

**Results:** We confirmed many known and identified 30 previously unreported candidate genes for milk, fat, and protein yield. While all of the genes were functionally associated with the traits, some showed pleiotropic effects as well. Specifically, association with mammary gland development, fatty acid synthesis, metabolism of lipids, or milk production QTLs in other farm animals has been reported. Variants associated with these genes explained a large percentage of genetic variance, compared to random ones.

**Conclusions:** Our findings proved the power of large samples and sequence-based GWAS in detecting new association signals. In order to fully exploit the power of GWAS, one should aim at very large samples combined with whole-genome sequence data. Although milk production traits in cattle are comprehensively researched, the genetic background of these traits is still not fully understood, with the potential for many new associations to be revealed, as shown in our study. With constantly growing sample sizes, we expect more insights into the genetic architecture of production traits in the future.

## Background

Intensive selection for milk production traits enhanced with improved nutrition and management, as well as reproductive technologies and accelerated by genomic selection (reviewed by [1]), has strongly increased milk production over the years [2]. The Holstein breed is dominant in milk production worldwide. The German Holstein population alone comprises 2.4 million cows, with an average milk yield of 10,000 kg per lactation [3]. The breeding goal for German Holstein is balanced and includes many traits that can be grouped into milk production, health, fertility, and longevity [4]. This has not always been the case, and although selection for milk production has been successful in increasing milk yield, it has also been associated with a higher incidence of mastitis, metabolic and reproductive diseases [5]. The relative weight of milk production in total merit indices is decreasing as new traits are continuously added to the breeding goal. However, because production still makes up a substantial part (e.g. 36% in Germany), there is the risk of a further decline in animal health. More extensive knowledge of the genetic architecture of economic traits is needed, especially given that the majority of these traits are complex traits, influenced by many genes and environmental factors.

So far, genome-wide association studies have been successful in the discovery of quantitative trait loci (QTL) and candidate genes (reviewed by [6]), however, only a few causal variants for economically important traits in cattle have been confirmed [7, 8]. In order to be able to detect the underlying causal variant whole-genome sequence (WGS) data and large samples are needed to ensure sufficient power of GWAS [9, 10]. GWAS in cattle is restricted by long-distance linkage disequilibrium (LD) segments [11], due to a small effective population size (N_e_) caused by intense selection [12], therefore making it hard to pinpoint the true causal variant which may be hidden among the many variants in LD. Another source of difficulty in revealing the true associations is the highly polygenic genetic architecture of quantitative traits, i.e. large number of variants with small effects affecting the trait [13]. Genotypes from whole-genome sequences obtained from sequencing the study individuals are limited, especially when large samples are considered. In that case, imputation [14] can be utilized as a method of obtaining the sequence-dense data. Imputation methods exploit LD patterns among the individuals in the sample and reference dataset, with the assumption that apparently unrelated animals inherited haplotype blocks from a common ancestor [15]. Imputation accuracy depends on various factors such as the size of the reference panel, the relationship between the individuals in the reference and sample dataset, imputation software choice and the number of the variants to be imputed [16–18]. In cattle, sequence-level imputation is usually performed in two steps [18], due to higher accuracy obtained when first imputing from a lower to a higher-density SNP chip, and then to sequence level.

To exploit the power of large sample size in detecting novel causative loci, we carried out GWAS for three milk production traits using imputed sequence data. After obtaining GWAS summary statistics with a mixed linear model approach, meta-analysis was utilized to pool the results of different animal groups. Candidate gene search was performed for top variants from GWAS with the lowest *p*-values and functional enrichment analysis was done to confirm the candidate genes. Finally, the percentage of genetic variance explained by the top SNPs was calculated to see which proportion of the variance could be attributed to variants associated with the novel candidate loci.

## Methods

### Dataset

The dataset for imputation consisted of 252,285 German Holstein cows with 45,613 SNP markers. Animals were mainly genotyped with various low-density SNP genotyping arrays (see Additional file 1: Table S1) and then imputed to 50K level according to the national genetic evaluation procedure [19], or genotyped with various 50K SNP chips (see Additional file 1: Table S1). The dataset was collected during the KuhVision project that aimed to genotype and phenotype German Holstein cows to establish a large-scale female reference population for genomic evaluation. The phenotypes for milk (MY), fat (FY), and protein yield (PY) in kg were obtained in the form of deregressed proofs (DRPs), which are pseudo-phenotypes produced using the special single-step SNP BLUP model for deregressing genomic estimated breeding values (GEBV) [20].

### Imputation

The genomic coordinates of the input genotypes were lifted from the previous bovine reference genome assembly UMD 3.1. to the ARS-UCD1.2 with a custom approach. CombineVariants from the Genome Analysis Toolkit (GATK) v. 3.8.1.0 [21] was used to merge the samples by chromosomes and by groups. The sample of 252,285 cows consisting of 30 autosomal and sex chromosome pairs was imputed to sequence level in a two-step imputation approach using the BEAGLE v. 5.2 [22]. The effective population size parameter was set to 1000. The animals were first imputed to high-density (HD) genotype level using the genotype data of 1278 Holstein cows consisting of 585,517 markers [23]. The HD reference panel was phased using BEAGLE v. 5.1 beforehand [24]. In the next step, data were imputed to the WGS level using the multi-breed reference panel from the 1000 Bulls Genome Project Run9 [25]. The reference panel consisted of 5116 cows and bulls of the species *Bos taurus* (see Additional file 1: Table S2). Both imputation steps were performed chromosome-wise, with the samples divided into groups of approximately equal size, due to high computational demand. The imputed files were indexed afterwards with IndexFeatureFile, GATK v. 4.2.2.0, merged by the sample groups, and split into separate lines due to the presence of multi-allelic variants (SNPs, insertions, and deletions) using BCFtools v. 1.14 [26]. As a quality control, the imputed WGS dataset was filtered using the dosage R-squared parameter, a measure of the estimated squared correlation between estimated and true allele dosage (DR2; [27]). Markers imputed with DR2 < 0.75 were removed with BCFtools. The imputed WGS dataset was annotated with VariantAnnotator from the GATK v. 4.2.2.0 using the Ensembl variation database, release 105 [28] imported from dbSNP [29], to account for SNPs without rsID.

### GWAS

Since phenotype measurements were not available for all 252,285 animals, the sample for GWAS consisted of 180,217 WGS-imputed cows with phenotypic observations for MY, FY, and PY. Samples were filtered for minor allele frequency (MAF) > 0.01. Due to memory restrictions of the high-performance computing (HPC) cluster, the samples were divided into 4 groups consisting of ∼ 45,000 animals each. GWAS was performed using the GCTA software v. 1.93.2 beta [30] applying a mixed linear model approach (MLMA) for all autosomes. The SNP effects were estimated using the following model:

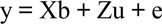

where y is a vector of DRPs; b is the fixed effect of the variant tested for the association with each trait; X is a vector containing the genotype score for the tested SNP; u is the vector of polygenic effects with u ∼ N (0, Gσ²u), where G is genomic relationship matrix (GRM) calculated using 50K SNP genotypes from all chromosomes, and σ²u is a variance of polygenic effects; Z is the incidence matrix of u; and e is the vector of residual effects with e ∼ N (0, Iσ²e), with I being an identity matrix and σ²e residual variance. Bonferroni correction was used to set a genome-wide significance threshold, corresponding to a *p*-value of 0.05/number of markers. The Manhattan plots were created using RStudio v. 3.6.3 [31]. METAL software [32] for meta-analysis was used to merge the GWAS summary statistics of each of the four animal groups per trait, using the approach that takes into account test statistics and standard errors. To correct for genomic inflation, lambda (λ_GC_) values were calculated as the median of observed χ² test statistics divided by the expected median of χ² distribution with one degree of freedom.

### Downstream analyses

SnpEff [33] and SnpSift [34] were utilized for functional annotation of genome-wide significant variants and prediction of their effect on genes, as well as the identification of the closest genes. The R packages cluster profiler [35] and DOSE [36] were used to carry out an over-representation analysis (ORA) [37] to determine whether the genes positioned closest to the genome-wide significant variants are enriched in the known biological pathways. ORA was performed using the Kyoto Encyclopedia of Genes and Genomes (KEGG) [38] database for variants that passed the significance threshold of 0.001/number of markers with enrichKEGG (*q*-value > 0.25). Candidate genes were also investigated manually, through the Animal Quantitative Trait Loci database (Animal QTLdb) [39] and using the publications previously associated with milk production candidate genes, based on the STRING database [40]. A Venn diagram of common candidate genes was created using the R package VennDiagram [41]. The percentage of genetic variance explained by the top 50 genome-wide significant SNPs and 50 random SNPs across all chromosomes was calculated using GCTA’s genomic-relatedness-based restricted maximum-likelihood (GREML) approach [42], by fitting the GRMs together in the model with 50K SNP chip variants. The analysis was done for the smaller subset of 45,000 animals due to high computational demand. PLINK v. 1.9 [43] was used to prune the variants in high linkage disequilibrium, based on pairwise *R²* correlation greater than 0.5 (--indep-pairwise 50 5 0.5).

## Results

### Imputation

To evaluate the genotype liftover quality, we examined the allele frequency (AF) concordance between the imputed WGS dataset and Run9, by plotting the AF from BTA16 of the two datasets against each other. Allele frequencies of imputed variants were congruent with the ones from 1000 Bulls Run9, showing the coherence in the frequency for the majority of loci (Figure S1). Imputation quality control was carried out by utilizing the DR2 parameter, built into the BEAGLE software. Markers imputed with DR2 < 0.75 were removed with BCFtools, leaving the 20,737,793 markers for further analyses. Then, we checked the DR2 values of known causal variants, such as two variants in the *DGAT1* gene, which were imputed with almost perfect quality (DR2=0.99), as well as rs385640152 in the *GHR* gene with DR2=0.98, and rs211210569 in *MGST1* with DR2=1. After the imputation of 252,285 animals to sequence level, and filtering for DR2 and MAF, 17,256,703 variants were left for GWAS.

### GWAS

A large number of variants exceeded the genome-wide significance threshold. GWAS analyses identified 54,032 significant variants for MY, 42,323 for FY, and 35,106 for PY, with the highest number of associations on chromosomes 5, 6, and 14 (Figures 1-3). Low *p*-values were observed for many SNPs, with top variants positioned on bovine chromosome (BTA) 14: rs109050667 (*p* = 7.04×10^-737^), rs136630297 (*p* = 7.18×10^-380^), and rs109050667 (*p* = 2.38×10^-221^) for MY, FY and PY, respectively

**Figure 1.**
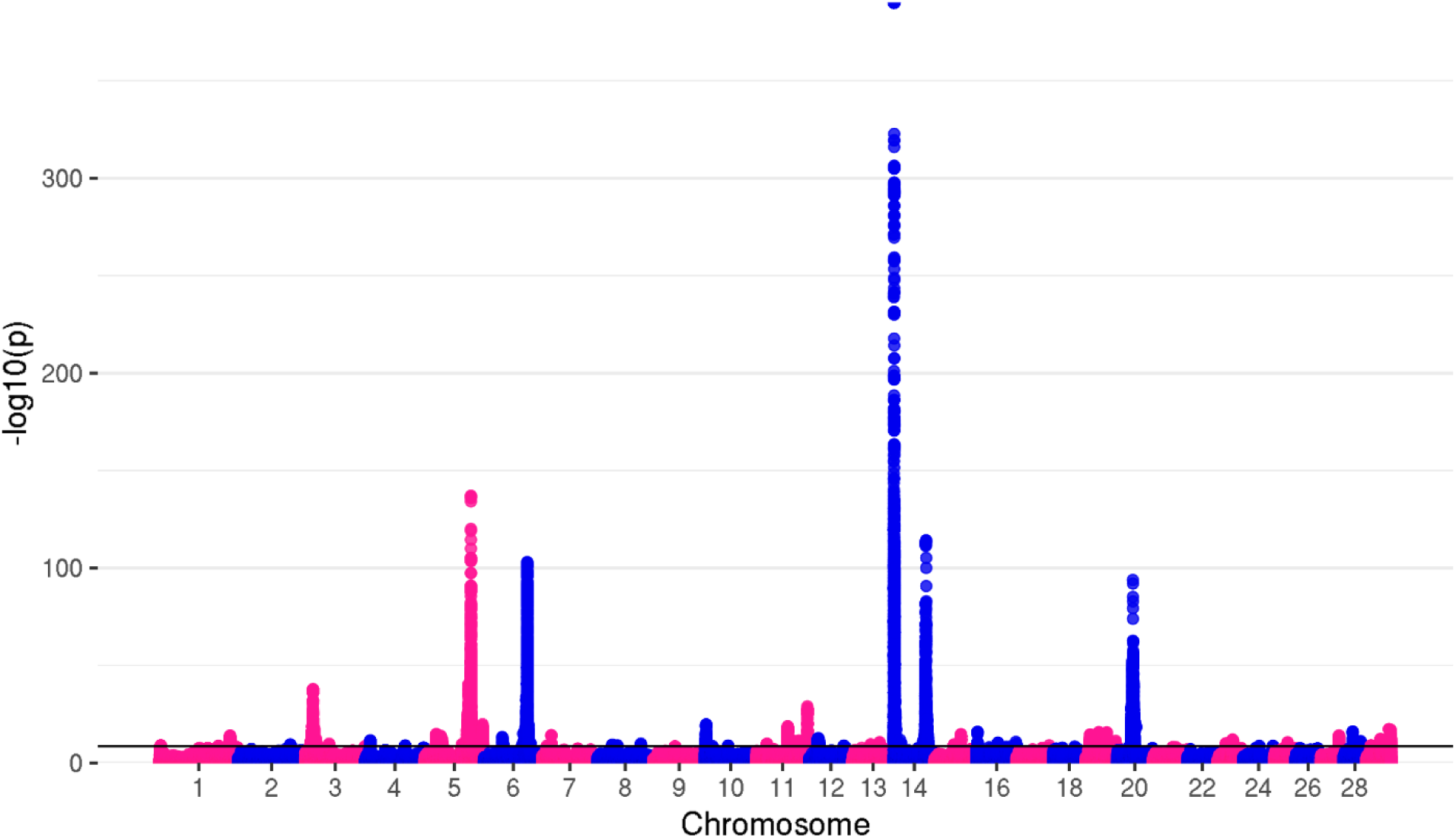
Manhattan plot for milk yield The top genome-wide SNP (*p* = 7.04×10^-737^) for MY was located on BTA14. However, RStudio used for the creation of this plot was not able to show *p*-values < 10×10^-325^, reporting them as “0”. Therefore, ylim had to be set lower, to provisional ylim of 390, in order to present all significant variants

**Figure 2.**
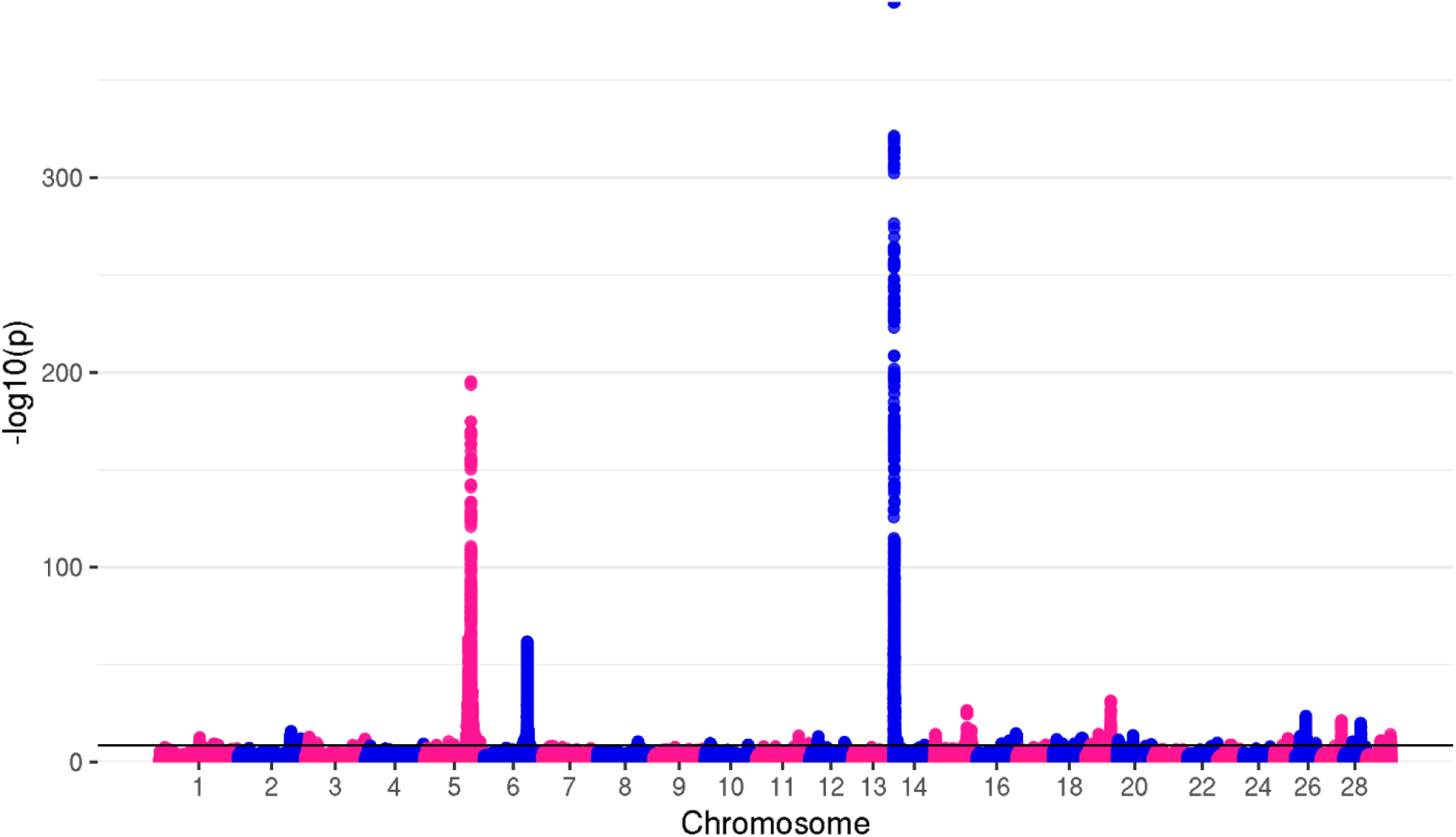
Manhattan plot for fat yield The top genome-wide SNP (*p* = 7.18×10^-380^) for FY was positioned on BTA14. However, RStudio used for the creation of this plot was not able to show *p*-values < 10×10^-325^, reporting them as “0”. Therefore, ylim had to be set lower, to provisional ylim of 390, in order to present all significant variants

**Figure 3.**
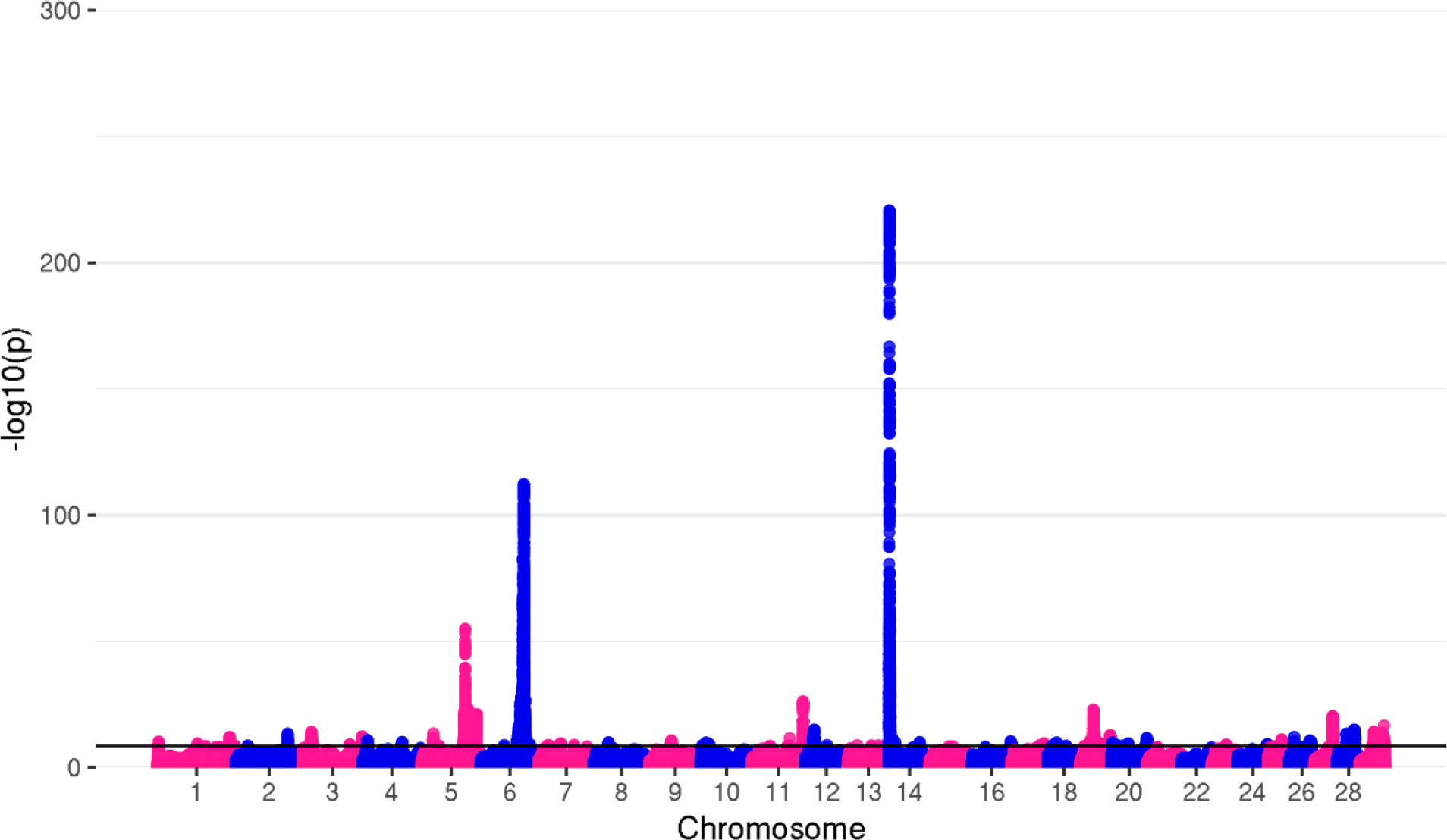
Manhattan plot for protein yield

The top 50 variants from each chromosome were chosen for further research (see Additional file 2: Tables S1-S3). Considering that significant associations have not been identified on every chromosome and that some chromosomes had less than 50 significant variants, the number of top variants chosen for further investigation differed across chromosomes and traits. For MY, 1012 top SNPs were found within or in proximity of 109 genes from 25 chromosomes (see Additional file 1: Table S3). The top candidate genes on chromosomes with the highest number of significant SNPs were *MGST1* and *SLC15A5* on BTA5, *GC*, *NPFFR2*, *ENSBTAG00000049290* and *SLC4A4* on BTA6, *ADCK5*, *CPSF1*, *SLC52A2*, *SLC39A4*, *FBXL6*, *TMEM249* and *SCRT1* on BTA14.

For FY, the top 962 SNPs from 24 chromosomes were located within or close to 108 genes (see Additional file 1: Table S3). The top candidate genes were *MGST1* and *SLC15A5* on BTA5, *GC*, *NPFFR2*, *ENSBTAG00000049290* on BTA6, *CPSF1*, *SLC39A4*, *ADCK5*, *TMEM249*, *SCRT1*, *SLC52A2*, *FBXL6* and *ENSBTAG00000053637* on BTA14.

For PY, 1065 top SNPs from 26 chromosomes were located close to or in 172 genes (see Additional file 1: Table S3). The candidate genes associated with the most significant genomic regions were: *GC*, *NPFFR2*, *ENSBTAG00000049290*, and *SLC4A4* on BTA6, *ABCC9*, *ST8SIA1*, *ENSBTAG00000026611* and *CMAS* on BTA5. Many genes were found to be associated with the same traits, as shown on the Venn diagram (Figure 4). The highest number of common candidate genes were found between MY and PY (47). The second highest number of candidate genes was between FY and PY (27), 7 genes were in common for MY and FY, and 23 genes were in common for all three traits (see Additional file 1: Table S4). Lambda values, calculated to assess for false associations were as follows: λ_MY_ = 1.764, λ_FY_ = 1.898, and λ_PY_ = 1.928. The reason for increased genomic inflation factors was due to the meta-analysis that inflated the *p*-values and therefore the number of genome-wide significant variants. To assess the effect of meta-analysis on inflation we divided the individuals from direct-GWAS summary statistics into smaller groups, running the GWAS for each of these groups again, and merging them into a meta-analysis. The lambda values were higher after merging the animals into meta-analysis compared to direct GWAS summary statistics for the same individuals (Figure 5).

**Figure 4.**
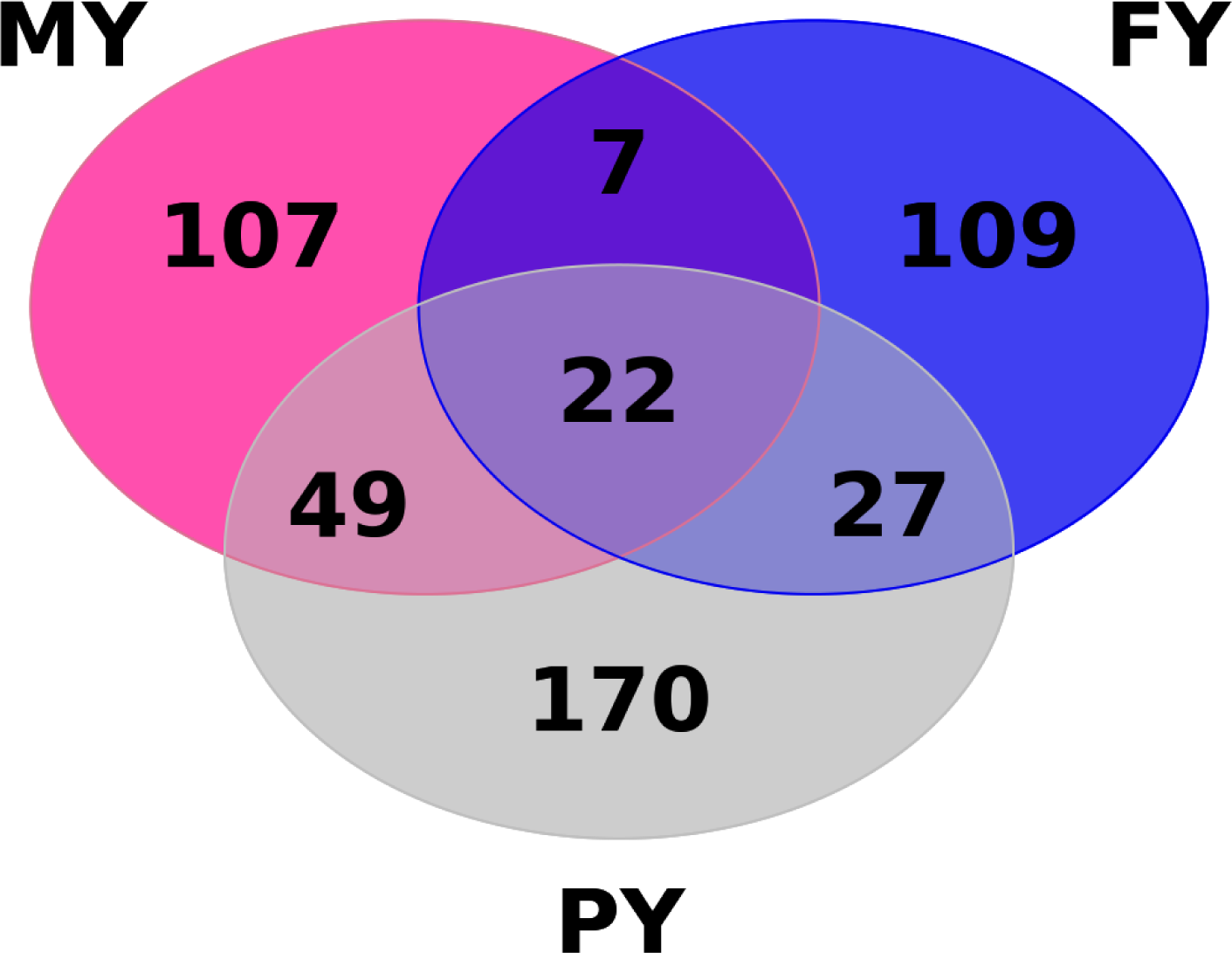
Venn diagram of MY, FY, and PY showing concordant and discordant candidate genes

**Figure 5.**
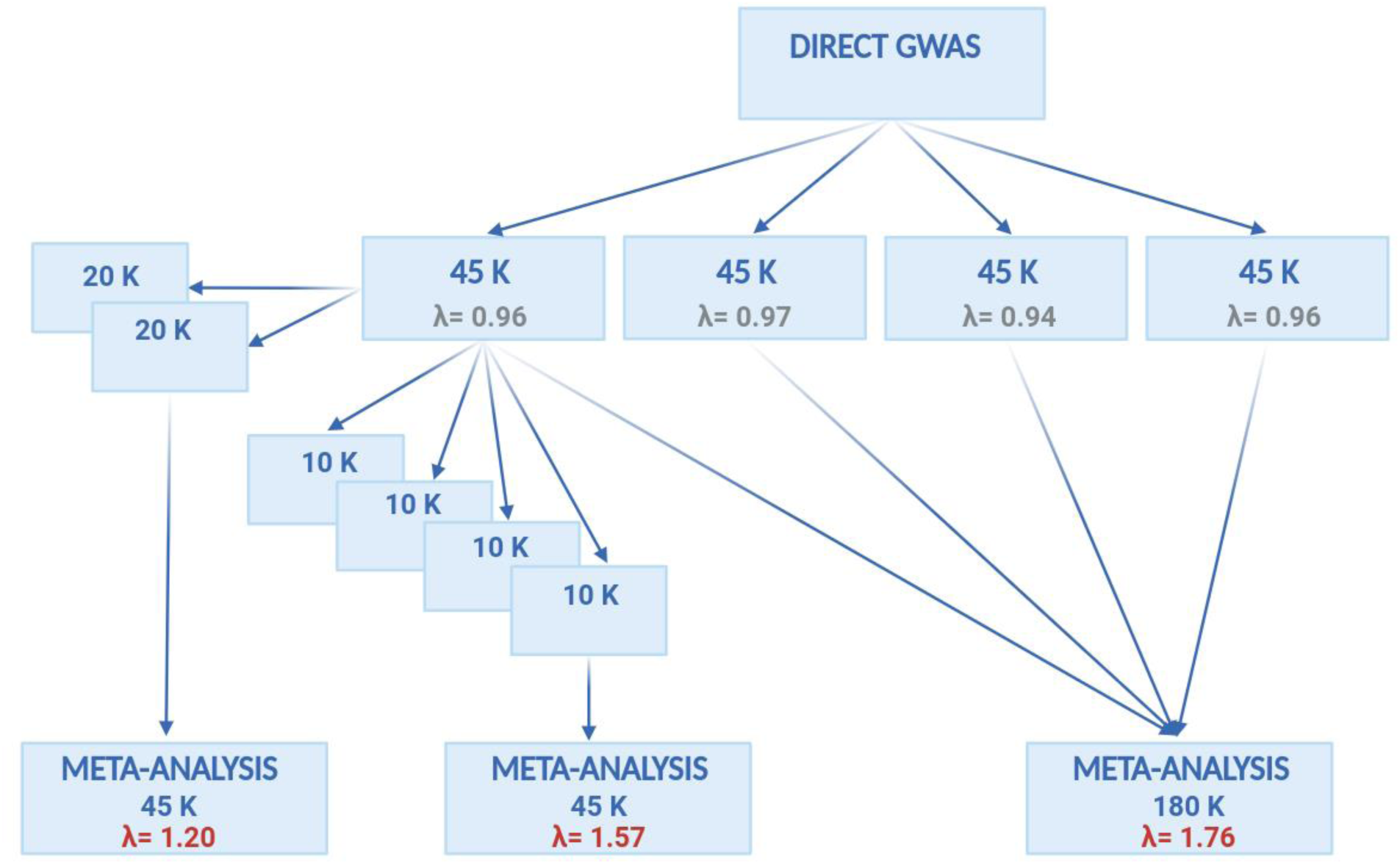
Genomic inflation factors of MY measured on direct GWAS summary statistics and after meta-analysis To check the cause of genomic inflation in meta-analysis summary statistics, one of the animal groups on which we ran direct GWAS was divided into two groups. For each of the two groups, GWAS was run again, and summary statistics were merged into the meta-analysis. Lambda values obtained on meta-analysis summary statistics were higher (λ= 1.20) than ones measured for the same individuals on direct GWAS summary statistics (λ= 0.96). To further check the extent of inflation caused by meta-analysis, the same group of animals was divided again, this time, into four groups. GWAS was run for each of the groups and results were merged into the meta-analysis. Lambda values were even higher this time (λ= 1.57). The figure was created with BioRender.com

### Downstream analyses

SnpEff was used to predict the functional effects of genome-wide significantly associated variants on proteins and to identify the closest genes. The majority of variants were identified in introns (46.41%) or intergenic regions (37.46%). The number of predicted effects was larger than the actual number of variants, due to genes with multiple transcripts and variants which affect multiple genes. A detailed description of the variant effects by type is available in an additional file (see Additional file 1: Table S5). Regarding the variant impact on proteins, a high majority of variants were classified as modifiers (98.38%), and only 0.025% were high-impact variants. Of the 50 top variants which were further investigated, the same high-impact, frameshift variant was found for both PY and MY on BTA16, at 80,129,589 bp, in the *SYT2* gene. One frameshift variant was also found for FY on BTA3, at 7,933,141 bp in the *FCGR2B* gene.

KEGG functional enrichment analysis revealed a large number of over-represented terms. To narrow the list of possible terms, ORA was performed only for genes associated with variants that passed the genome-wide significance threshold of 0.001/number of markers. A list of all over-represented genes and associated pathways is available in an additional file (see Additional file 1: Table S6). The common dot plot of the 20 most significant terms of KEGG analysis for MY, FY, and PY is shown in Figure 6. The top variants were found in or in proximity to the genes over-represented in 23 pathways, mostly in the PI3K-Akt signaling pathway (Table 1). Two terms were significantly enriched with all three traits.

**Figure 6.**
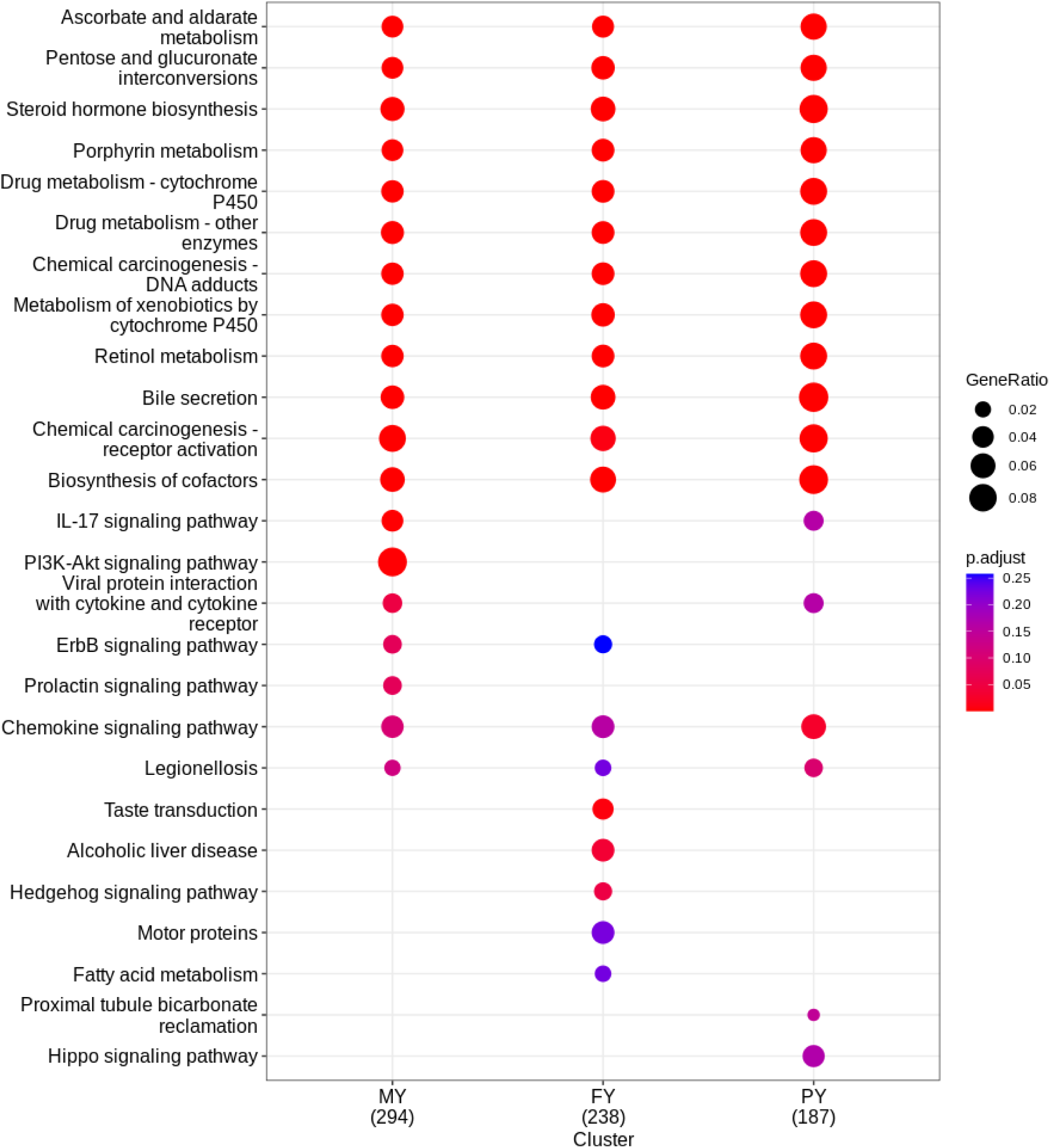
Enrich KEGG dot plot of 18 most significant pathways for MY, FY, and PY

**Table 1.**
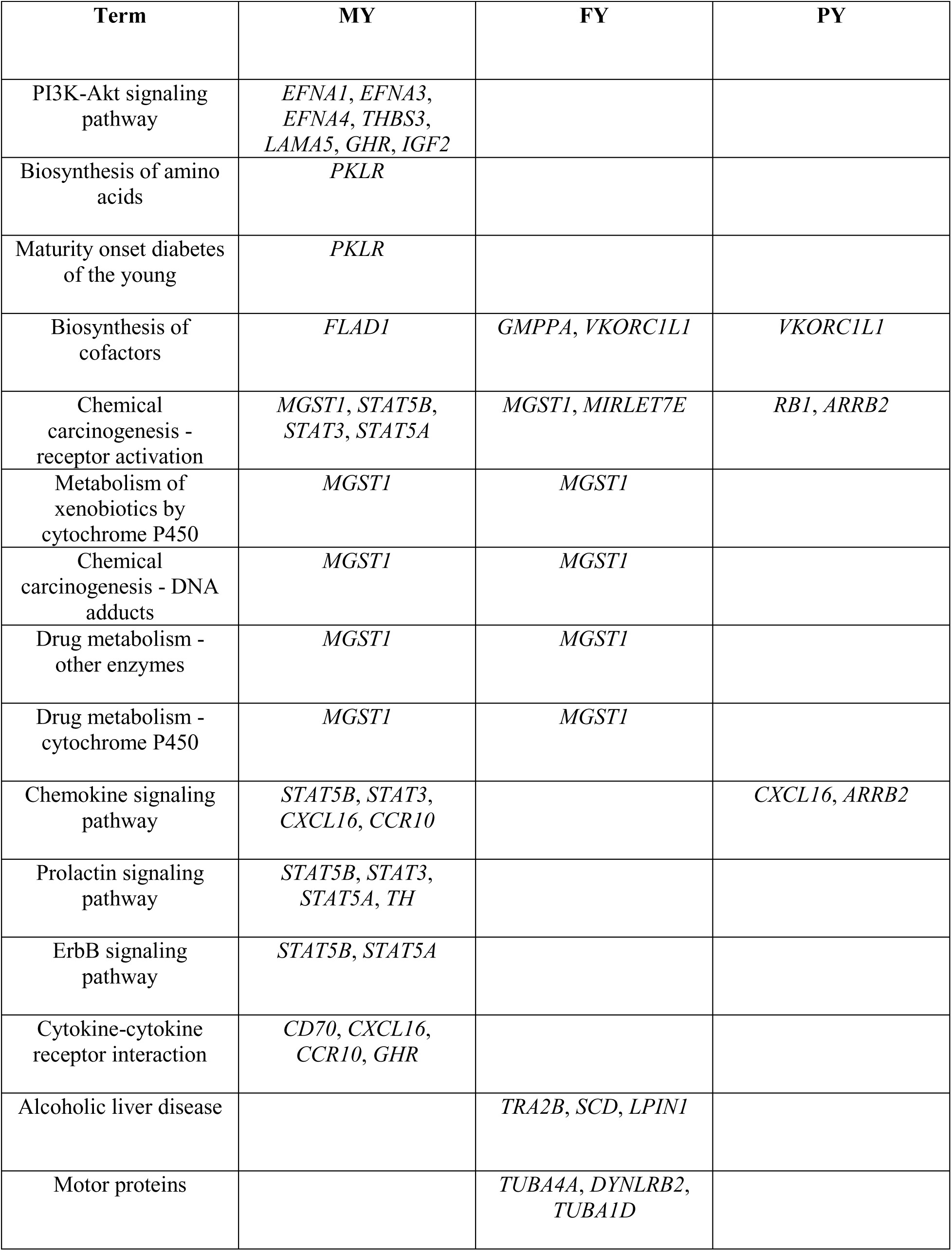
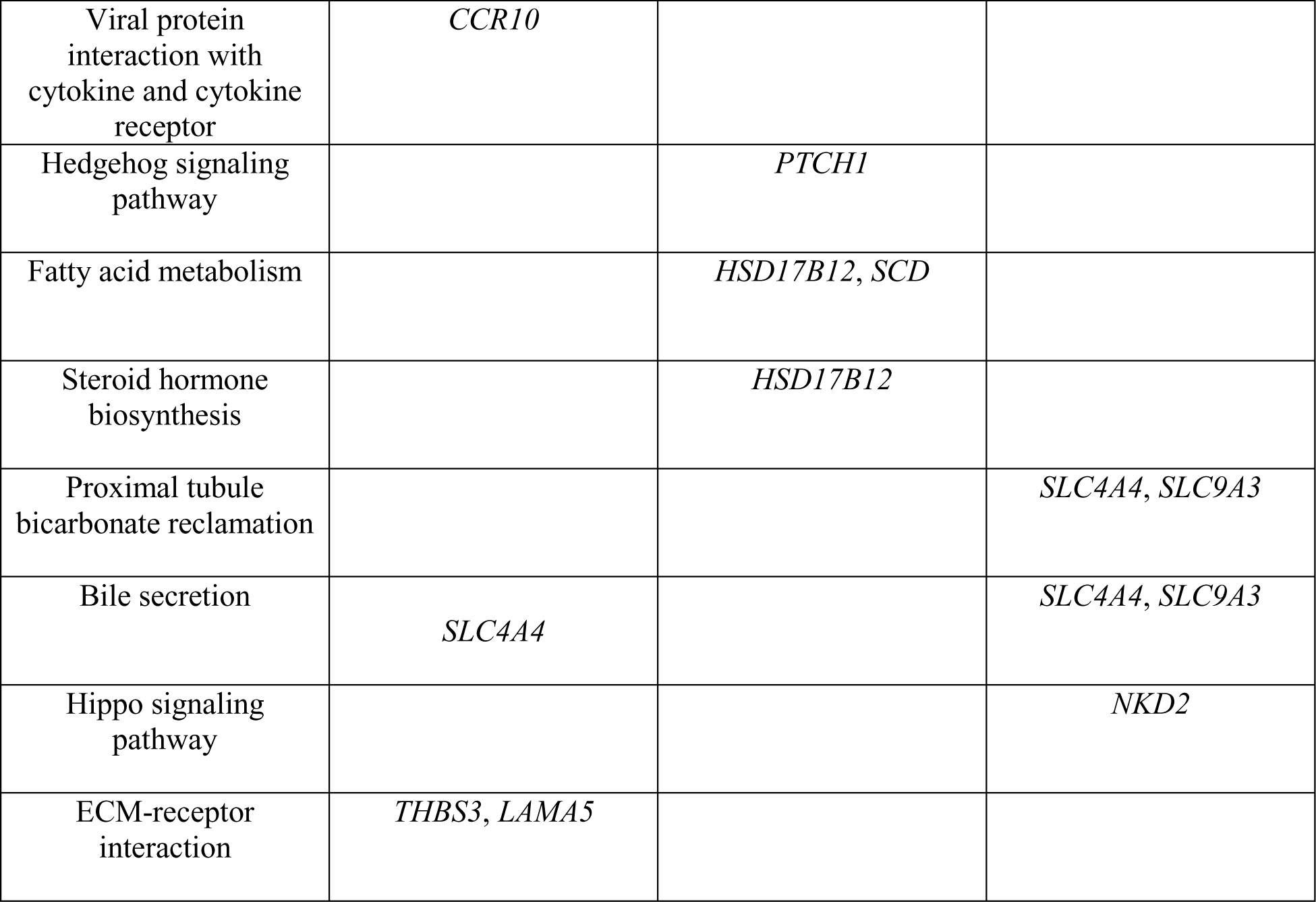
Over-represented genes associated with top variants for MY, FY, and PY.

The percentage of genetic variance explained by 50 top variants, as well as by 50 random variants from all autosomal chromosomes was estimated for all three traits (Table 2). For MY, 1012 top variants from 25 chromosomes explained 8.67% of the variance. Random SNPs from 29 autosomal chromosomes, explained on average 0.78% of the variance. As for FY, 962 top SNPs from 24 chromosomes accounted for 7.04% of the genetic variance, while the random 1450 variants from all chromosomes explained about 0.31%. For PY, 6.66% of the variance was explained by 1065 top SNPs from 26 chromosomes, and 0.37% was due to random 1450 variants. After LD pruning of the top variants for each trait, there were 124, 147, and 182 variants left for MY, FY, and PY, respectively. Genetic variance explained by pruned variants was 10.01% for MY, 6.51% for FY, and 5.17% for PY.

**Table 2.** Genetic variance explained by top and random variants for MY, FY, and PY.

| **Trait** | **V<sub>TOP</sub>** | **SE<sub>TOP</sub>** | **V<sub>RANDOM</sub>** | **SE<sub>RANDOM</sub>** |
| --- | --- | --- | --- | --- |
| **MY** | 0.086677 | 0.012530 | 0.003195 | 0.002656 |
|  |  |  | 0.015675 | 0.003461 |
|  |  |  | 0.005690 | 0.002973 |
|  |  |  | 0.005522 | 0.002872 |
|  |  |  | 0.008892 | 0.003040 |
| **FY** | 0.070413 | 0.010478 | 0.003675 | 0.002497 |
|  |  |  | 0.001819 | 0.002540 |
|  |  |  | 0.004907 | 0.002939 |
|  |  |  | 0.001645 | 0.002567 |
|  |  |  | 0.003471 | 0.002572 |
| **PY** | 0.066613 | 0.009325 | 0.003456 | 0.002645 |
|  |  |  | 0.007485 | 0.002946 |
|  |  |  | 0.003309 | 0.002646 |
|  |  |  | 0.001254 | 0.002492 |
|  |  |  | 0.002862 | 0.002649 |

$V_{\text{TOP}}$ = genetic variance explained by top genome-wide significant variants from autosomal chromosomes
$SE_{\text{TOP}}$ = standard error of top variants
$V_{\text{RANDOM}}$ = genetic variance explained by random 50 variants from all autosomal chromosome
$SE_{\text{RANDOM}}$ = standard error of random variants

## Discussion

### Imputation

In this study, we performed a stepwise imputation of 252,285 German Holstein cows from SNP chip up to sequence level, which makes our sample size one of the largest imputed in cattle so far. The stepwise imputation approach seems to improve the imputation accuracy, as previously shown in cattle [18, 44]. Imputation error rate tends to decrease when an intermediate reference panel is used [44], possibly due to a larger choice of possible haplotype matches between WGS and medium-density SNP chip, which are narrowed down when using an HD panel as an intermediate [18]. In our study, stepwise imputation was done using the Holstein breed HD panel, a subset from van den Berg et al. [23] as an intermediate step, and the WGS panel from 1000 Bulls Genome Consortium, as a second step. The WGS-based panel consisted of various breeds of taurine cattle (see Additional file 1: Table S2). The usage of a multi-breed reference was shown to increase the imputation accuracy in many studies in cows [45–47], especially for low-frequency variants [46]. However, multi-breed panels can be counter-productive if animals in the reference panel are too distant from the sample dataset [48]. The usage of BEAGLE software for imputation can at least partly overcome this issue since its algorithms can prioritize between the closer and the genetically distant individuals in the multi-breed reference panel [49]. Moreover, the 1000 Bulls reference panel consisted of a large number of Holstein animals (∼1200) making them the most represented breed in the reference panel (see Additional file 1: Table S2), therefore enabling the reliable imputation of Holsteins even in the presence of genetically distant breeds. Another crucial factor to consider is the value of the N_e_ parameter [49]. Default Ne in BEAGLE is 1,000,000, however, this corresponds to the human populations for which was it initially developed. Therefore, updating the Ne parameter to smaller values is needed, when working with other, less-diverse populations [49].

To evaluate the accuracy of imputation we used the second category of quality measures [50] based on estimated genotypes (DR2) since SNP array genotyped animals were not whole genome sequenced. Filtering the variants with DR2 < 0.8 is recommended when using DR2, as the imputation error rate increases below this threshold [49] hence we filtered out all the variants with DR2 < 0.75. Known causal variants were left in the dataset after DR2 filtering, and were imputed with near to perfect quality (DR2=0.98 to 1). *DGAT1* causal variants were among the 100 top genome-wide significant variants for all three traits analyzed but were not the top variants. A possible explanation for this could be the presence of additional variation in the form of a known variable number of tandem repeats (VNTR) in the *DGAT1* region or low imputation accuracy [47, 51, 52]. To assess the liftover quality, AF concordance between the imputed WGS dataset and the Run9 reference panel was examined on the example of chromosome 16, showing the congruence for the majority of variants between the two datasets (see Additional file 1: Figure S1).

### GWAS and candidate gene research

After carrying out the GWAS, possible candidate genes were retrieved by searching public databases such as Animal QTLdb and reviewing journal papers on previously reported candidate genes and QTLs. We confirmed many of the previously reported QTLs and candidate genes (see Additional file 1: Table S7 and Additional file 3: Tables S1-S3) such as *DGAT1* and its variants rs109326954 and rs109234250 on BTA14, *MGST1* on BTA5, *ABCG2* on BTA6, *GC* on BTA6 and *GHR* on BTA20, but also discovered new, previously unreported loci (Table 3). There were a large number of genes whose functions have not been reported yet, as well as the ones whose functions could not be associated with milk production or content (see Additional File 4: Table S1). Therefore, these genes were not considered as potential candidate genes. For simplification, we discussed only candidate genes associated with the most significant SNPs, while the list of all associations can be found in Additional file 3: Tables S1-S3. The majority of the most significant variants were intronic (37%) and intergenic (30%) (see Additional file 1: Table S5). Most of the significant variants that were included in candidate gene research were non-coding as well, which is in line with the majority of other GWAS publications [53–55]. Nayeri et al. [55] showed that a large proportion of the most significant variants affecting milk yield and composition traits in Holstein and Jersey cattle were located in non-coding regions of the genome. Both intron and intergenic variants usually do not code for proteins, making their functional prediction challenging [56]. However, recent research in human studies (reviewed by [53]) and cattle [57] has shown that even the variants in non-coding regions may play an important part in complex traits and diseases, by indirect involvement in gene expression regulation. Known QTNs in livestock are not all coding variants that cause a change in amino acid [6, 58], therefore, variants in non-coding regions can be causal as well [57]. Xiang et al. [59] showed that non-coding variants can contribute substantially to variance in complex traits in cattle.

**Table 3.**
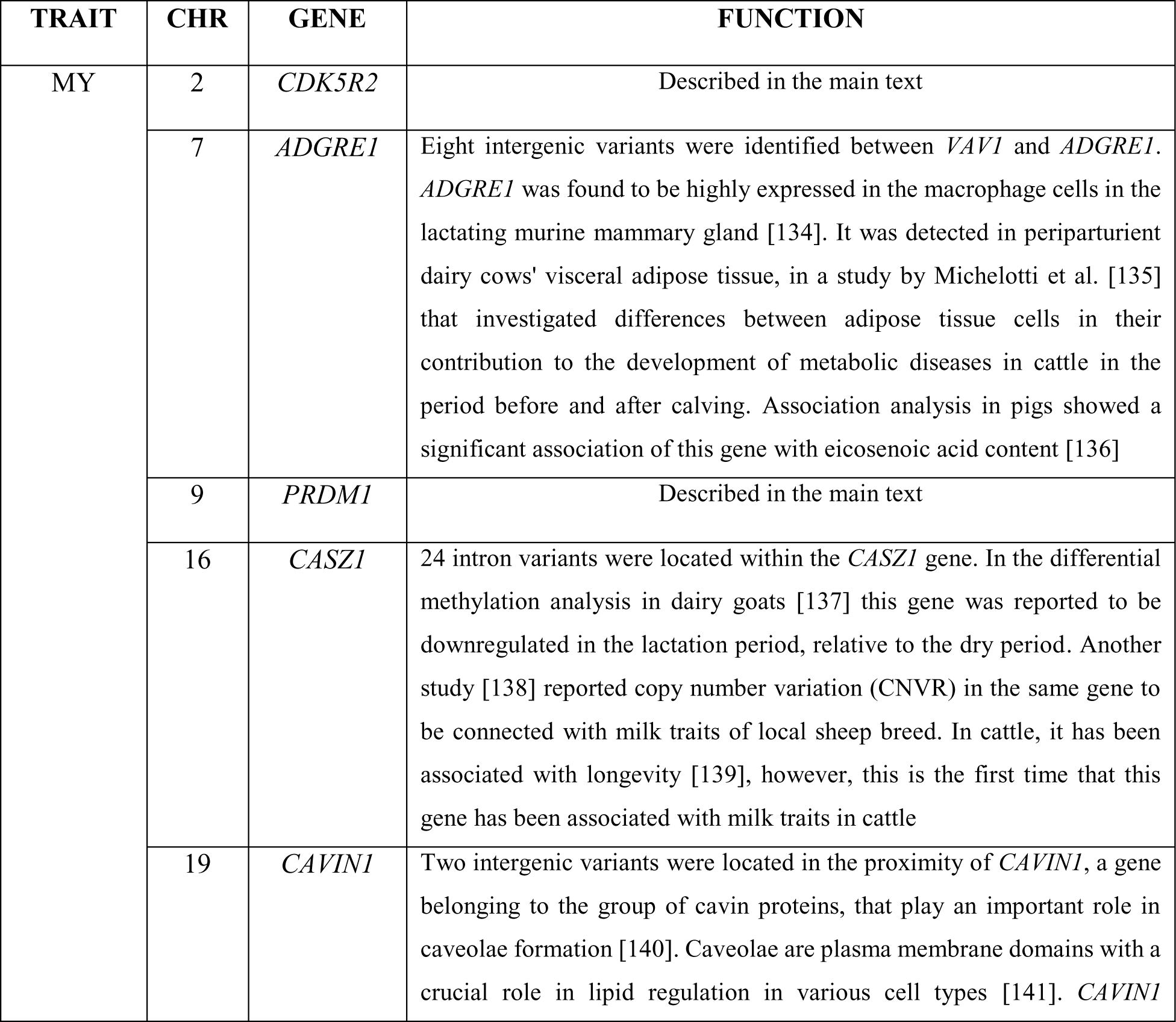
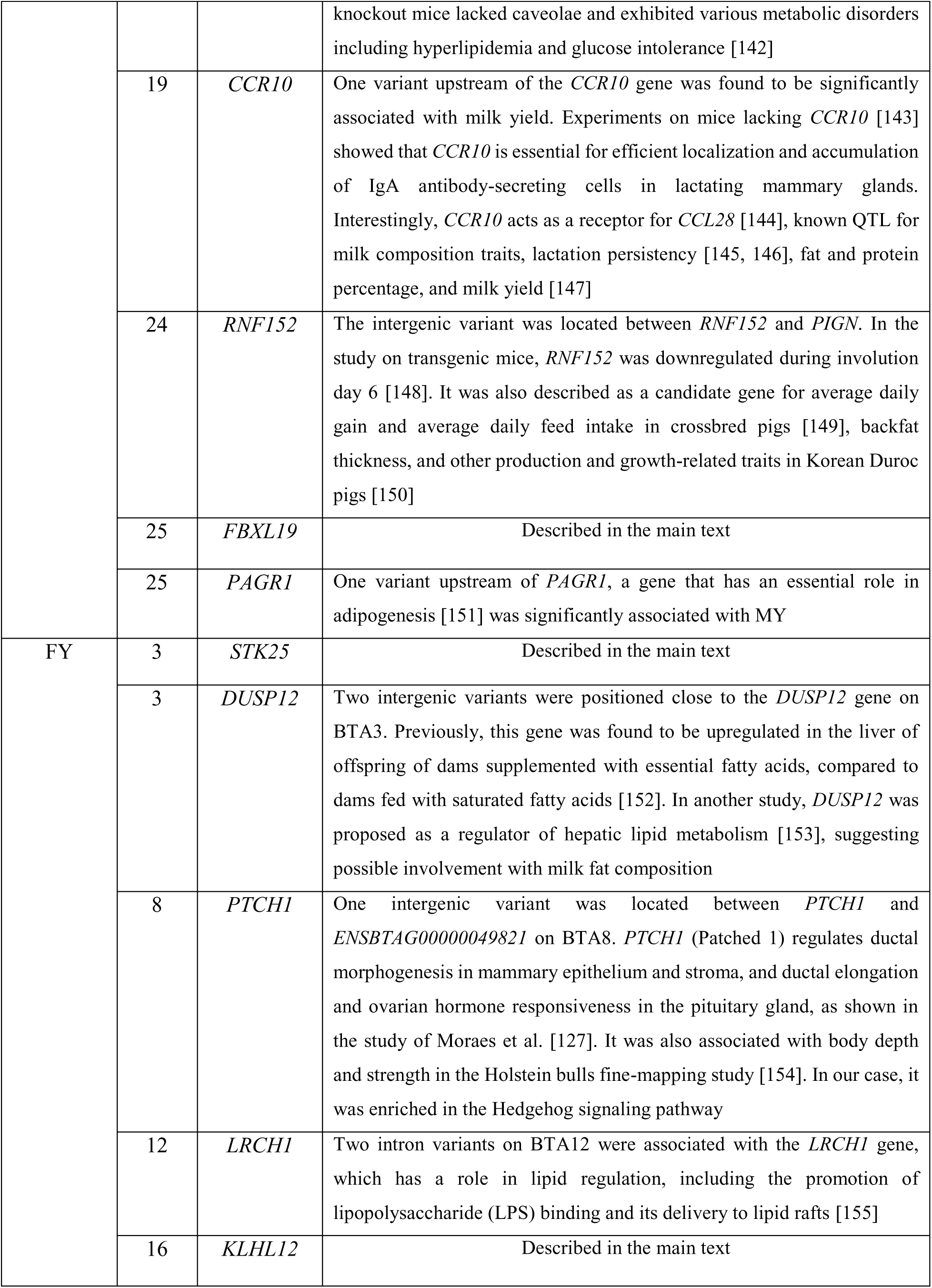
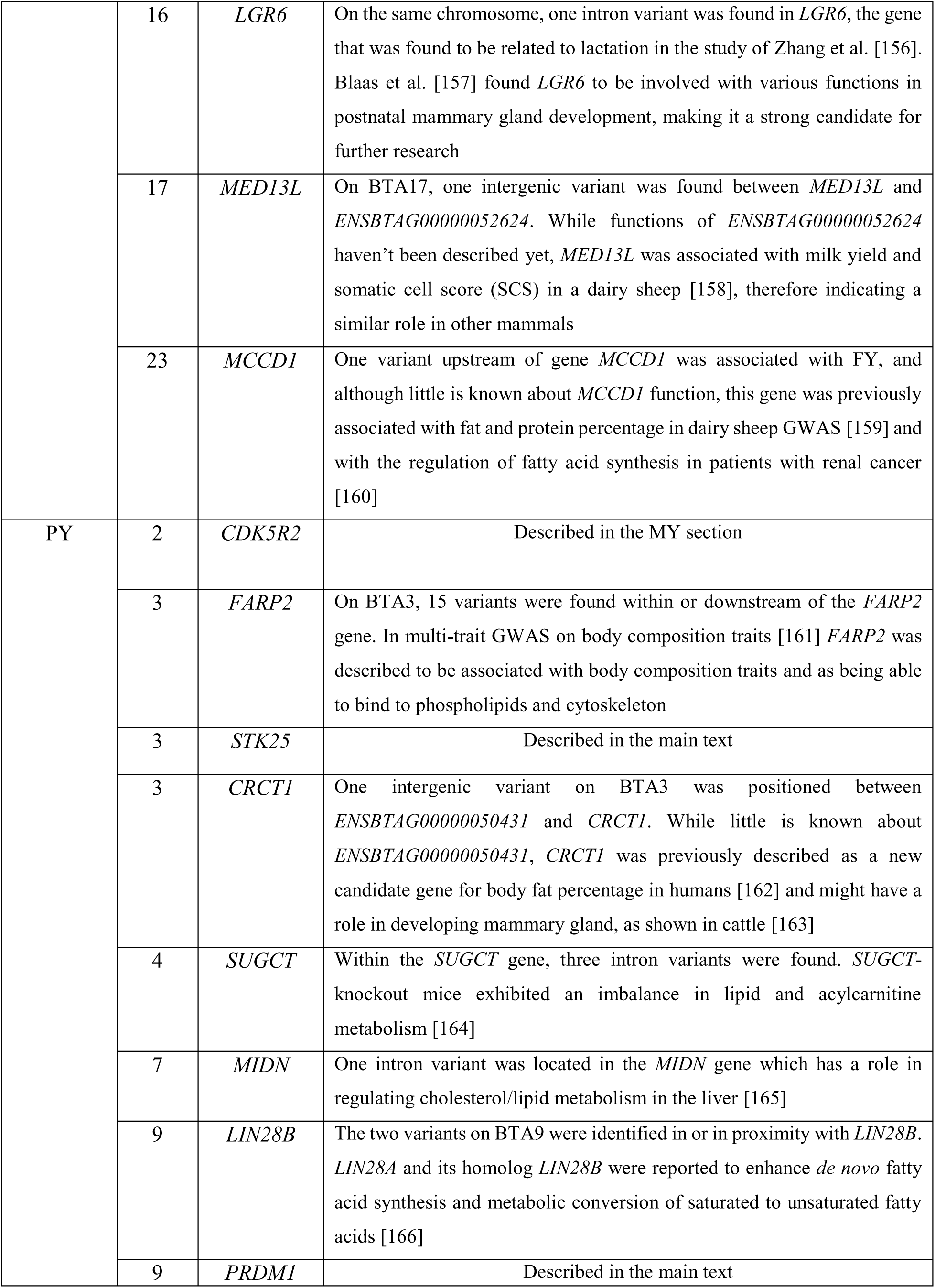
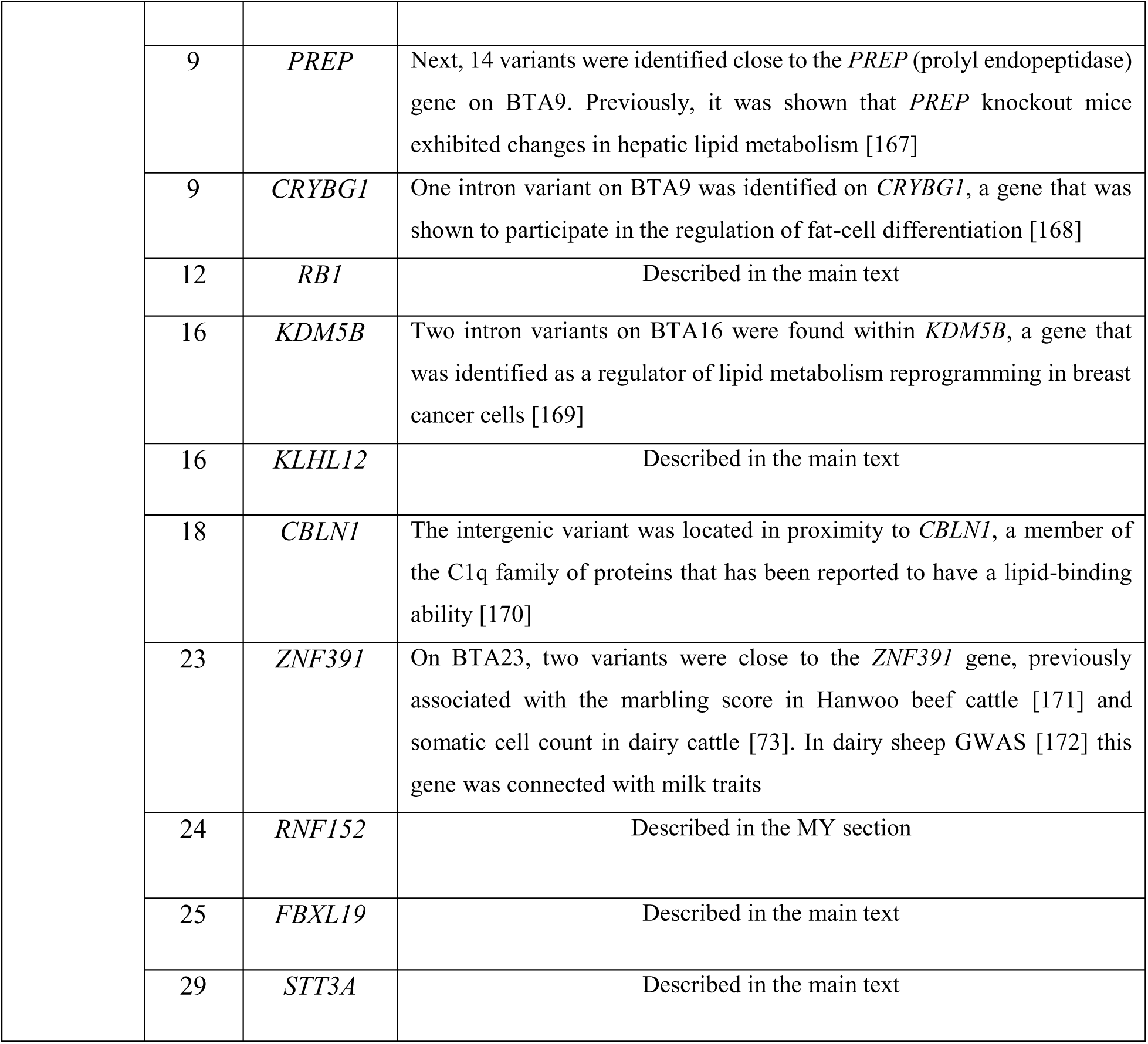
New candidate genes for milk production traits.

Due to the large sample sizes in our study, which might contribute to the rise in genomic inflation [60], lambda values were measured before and after performing the meta-analysis. Genomic inflation is a spurious association between a variant and trait, where the relationship between a phenotype and SNP seems to arise from different factors than the true association [61]. These factors include population stratification [62], cryptic relatedness [63], polygenic inheritance [61], or strong association between variant and phenotype [64]. Although some of the genomic inflation in our study might be attributed to the polygenicity of milk production traits [65], and population structure in German Holstein [66], the main source of genomic inflation was the use of meta-analysis software (Figure 5). Similar findings were reported in human studies [67], where large number of individuals are often pooled into the meta-analysis. The use of meta-analysis was inevitable in our case, due to the large samples that our HPC cluster was not able to utilize. MLMA accounted properly for genomic inflation, as the direct GWAS summary statistic had lambda values below 1 (Figure 5), and values up to 1 are usually considered as a threshold for genomic inflation. To prove that inflation was not due to population structure amplification that might arise when pooling the samples into the meta-analysis [68], we divided one of the animal groups on which we obtained summary statistics. After the animals were divided into two groups, GWAS was run for each of them again. Then, after obtaining the summary statistics, two groups of samples are merged into the meta-analysis. As shown in Figure 5, lambda values for the same samples were increased after pooling into a meta-analysis. Moreover, an increase in the number of animal groups pooled into meta-analysis led to higher genomic inflation (Figure 5). Considering that many factors that could lead to an increase in lambda values were present in our study, including the polygenic nature of the milk production traits, large sample sizes, potential underestimated relationships between animals, and in the end, the use of meta-analysis, we consider the values we obtained on meta-analyzed traits (1.764-1.928) acceptable even though they exceeded the generally accepted threshold of 1.

### Candidate genes for milk yield

The novel candidates that appeared to be the most relevant for further validation experiments due to their role in organism will be discussed here, while the list of all novel candidate genes and their roles connected with milk production traits are listed in Table 3. Except for the functional involvement of the candidate genes with milk production traits, and the fact that some of them are reported in other mammal species for the same or similar traits, variants found in candidate genes need experimental validation to be considered causative. For this purpose, prioritization of genome-wide significant variants according to external evolutionary and functional information [59] is suggested as the next step, followed by sequencing and gene editing experiments.

As for the new associations, we identified 9 genes that, to our knowledge, were not previously described in cattle for milk yield or related traits. On chromosome 2, we identified two intergenic variants whose positions fall between the *FEV* and *CDK5R2* genes. While *FEV* was reported earlier [69], *CDK5R2* has not been associated with milk traits in cattle yet. *CDK5R2* (Cyclin Dependent Kinase 5 Regulatory Subunit 2 (p39)) acts as one the activators of the *CDK5* gene [70] that has numerous important roles in the nervous system [71]. Talouarn et al. [72] identified the variants in the region of this gene to be associated with milk yield in French dairy goats. *CDK5R2* was previously associated with somatic cell count (SCC) in dairy cattle [73], and meat color traits in Nellore cattle [74], and crossbred and purebred pigs [75]. The variant in this gene has been associated with teat length in Chinese Holstein, in the study of Wu et al. [76]. Given the previous association with milk yield in the goat population, as well as with udder-related traits in cattle, this gene presents a strong candidate for further research.

A downstream variant of the *PRDM1* gene on BTA9 at 43,842,866 bp, was significantly associated with MY. *PRDM1* (PR Domain Zinc Finger Protein 1), or *BLIMP1* (B-Lymphocyte-Induced Maturation Protein 1) was described as an essential factor for mammary development in mouse studies [77]. Ahmed et al. [77] discovered that a group of luminal alveolar progenitor cells, expressing *BLIMP1*, were essential for mammary gland development. *BLIMP1* is required for ductal morphogenesis in puberty and alveolar maturation in pregnancy and lactation, with its inactivation causing inadequate milk secretion [77]. In another study [78] *BLIMP1* was described as necessary for the delay of the intestinal epithelium maturation from suckling to adult-type intestinal epithelium, with mutant mice showing growth disturbances and increased mortality.

One intron variant in the *FBXL19* gene was associated with MY. *FBXL19* (F-Box And Leucine Rich Repeat Protein 19) regulates cell migration and proliferation [79, 80] and acts as a major regulator of adipogenesis [81]. Adipose tissue is a source of energy for various organs and tissues, as well as for the mammary gland especially during lactation when it serves as a source for fatty acids synthesis [82]. Adipogenesis is essential for the efficiency of milk production in dairy cows, as well as reproduction [83] making this gene an interesting candidate for milk production traits.

### Candidate genes for fat yield

Eight novel candidate genes were associated with FY in our study. The majority were involved with various lipid metabolism functions, therefore, we will describe only a few in the main text, while the description of the rest of the genes and their functions is available in Table 3.

One intergenic region between the *STK25* and *BOK* was significantly associated with fat yield on BTA3. While *BOK*, a member of the family of BCL-2 proteins which are involved in many cellular processes [84], couldn’t be linked to milk production traits, *STK25* attracted our attention as a candidate for fat yield and composition. *STK25* (serine/threonine protein kinase 25) belongs to the germinal center kinase III subfamily of Ste20 (sterile 20) proteins that exhibit various cell functions (reviewed in [85]). *STK25* was shown to regulate lipid catabolism in liver cells in humans and the release of non-esterified fatty acids (NEFA) from lipid droplets [86]. High levels of NEFA seem to stimulate the expression of the *CIDEA* gene, and consequently increase fatty acid synthesis *de novo* and milk fat secretion [87]. *CIDEA* was recently described as a regulator of *de novo* fatty acid synthesis in cattle as well [88]. In general, high levels of NEFA are associated with ketosis and fatty liver, poor reproductive performance, and negative energy balance in early lactation (reviewed by [89]). Another study indicated a plausible role of this gene in the regulation of lipid and glucose metabolism in the skeletal muscle of rodents and humans [90]. Altogether, *STK25* seems to have an important role in lipid metabolism and therefore is recommended for further investigation.

On BTA16, one intron variant was found in *KLHL12*, a gene described as essential for the secretion of apolipoprotein B100 (apoB100) very low-density lipoprotein (VLDL) particles in rat hepatoma cell line [91]. ApoB100, a major component of VLDL, is essential for the transport of triglycerides, the main component of milk fat, from the liver to peripheral tissues [92, 93]. Decreased levels of apoB100 in cattle have been reported in cows with metabolic disorders such as ketosis, milk fever, and displaced abomasum [94, 95]. *KLHL12* (Kelch-like Family Member 12) is a member of the Kelch-like family (KLHL) of proteins with important functions in the ubiquitination of proteins as reviewed by Shi et al. [96]. When it comes to known roles of the *KLHL12*, it has been reported as a negative regulator of the Wnt signaling pathway [97], important for various cell functions in both adult and embryonal tissue homeostasis (reviewed in [98]). It also seems to have a key role in collagen secretion [99]. Everything considered, this gene could potentially affect not only milk production traits due to its role in triglyceride transport but health traits as well, and because of this further validation is needed.

### Candidate genes for protein yield

For protein yield, many known candidate genes, as well as the pleiotropic effects of some genes were confirmed (see Additional file 1: Table S7), while 18 genes from 12 chromosomes are reported here, for the first time (Table 3). As for the genes with pleiotropic effects, three intergenic variants were positioned between the *FEV* and *CDK5R2* on BTA2, a gene that we found to be a novel candidate for MY in the previous paragraph. The five variants on BTA3 were located in or downstream of the *STK25* gene, showing the effects of this new candidate gene on both fat and protein yield. Then, on BTA9, 21 variants were located in or in proximity to *PRDM1*, identified in both MY and PY GWAS. The majority of *PRDM1*-associated variants were intergenic, however, variant rs136669229 (*p* = 1.009×10^-10^) was identified as missense, causing the Valine to Phenylalanine amino acid change, however, there was no difference in protein structure prediction. On BTA16, one intron variant was located within the *KLHL12* gene, whose function is described in detail in the FY section. Finally, we identified an intron variant in the *FBXL19* gene, our candidate gene for MY.

In the proximity of *STT3A* on BTA9, one variant was significantly associated with PY. *STT3A* (STT3 Oligosaccharyltransferase Complex Catalytic Subunit A) encodes the protein which is a part of the central enzyme complex in glycosylation [84]. Glycosylation is a post-translational protein modification that takes place in the endoplasmic reticulum (ER) and is essential for numerous cellular functions [100]. The two main types of glycosylation are N and O-glycosylation. N-glycosylation consists of the attachment of an oligosaccharide N-Acetylglucosamine to Asparagine residues and occurs in both eukaryotes and prokaryotes [101, 102]. The most important step in N-glycosylation is catalyzed by the oligosaccharyltransferase (OST) complex, consisting of different subunits of which the STT3 subunit is the most important [103]. In the study of human milk lactoferrin glycosylation [104], the expression of *STT3* in milk somatic cells decreased from day 4 to day 15 of lactation, leading to changes in the overall level of glycosylation [104]. Lactoferrin is a milk-derived glycoprotein with many important roles in organism including immunomodulatory and anti-inflammatory, anticancer, and antimicrobial functions [105]. Therefore, the *STT3A* gene might affect the protein yield, and possibly play a role in mastitis given the antibacterial function [105] of lactoferrin.

An intron variant was found within the *RB1* (retinoblastoma 1) gene on BTA12. *RB1* is known for its role in regulating the metabolism of glycolipids in the liver, muscle, and adipose tissues and improving fat and protein metabolism disorders [106]. A study on *RB1* knockout-mouse showed the potential involvement of *RB1* in the gut microbiota and intestinal free fatty acids profiles [107], altogether making this gene a strong candidate for further research.

### Downstream analyses

In the KEGG enrichment analysis, a large number of trait-associated genes were found within various pathways and biological processes. However, we restricted our analysis only to genes containing one of the top SNPs (Table 3). The highest number of genes (7) were involved in the PI3K-Akt signaling pathway, one of the most important signaling pathways that affect many biological functions, including cell metabolism, growth, migration, proliferation, and survival [108, 109]. Hou et al. [110] showed that *EEF1D* regulates milk lipid secretion and mammary gland development through interaction with various pathways, including PI3K-Akt. Genes involved in this pathway (Table 1) were all previously reported as candidates for milk production and composition traits (see Additional file 1: Table S7). Other pathways and terms involved with milk composition, synthesis, and secretion or mammary development processes included biosynthesis of amino acids, biosynthesis of cofactors, prolactin signaling pathway [111], ErbB signaling pathway [112], Hedgehog signaling pathway [113], fatty acid metabolism, Hippo signaling pathway [114] and ECM-receptor interaction [115]. The term “biosynthesis of amino acids” included the gene *PKLR*, a known candidate gene for milk yield and composition traits [116], with a role in the regulation of triglyceride levels and fatty acid synthesis [117]. KEGG category “biosynthesis of cofactors” contained three genes, across the three traits. Of these genes, *FLAD1* was associated with MY in our study and was previously reported as a candidate gene for milk calcium content and lactose percentage [118, 119]. Expression of gene *GMPPA*, whose variants were associated with FY, was positively correlated with bovine milk fat globule size in the study of Huang et al. [120]. This pathway was also enriched with *VKORC1L1,* a gene involved in vitamin K metabolism [121], across two traits. Although this gene couldn’t be linked to milk production, it was described as a candidate gene for subclinical ketosis in Holstein in the study of Soares et al. [122]. Interestingly, it has an important role in adipogenesis, with *VKORC1L1* mutated mice having smaller length and weight than wild type [123], making the possible role in milk fat metabolism plausible. Prolactin signaling pathway was enriched with genes *TH*, *STAT5A*, *STAT5B*, and *STAT3* for MY. Prolactin (*PRL*) is a gene well-known for its role in mammary development and lactation in many mammal species, as well as in cattle [111, 124]. *STAT5A*, *STAT5B*, and *STAT3* genes belong to the STAT family of transcription factors that participate in the PRL receptor (*PRLR*) signaling pathway [111] and were previously associated with milk composition traits in GWAS in Holstein cattle [125]. *STAT5A* and *STAT5B* were also enriched in the ErbB signaling pathway for MY. The members of the ErbB family of tyrosine kinase receptors regulate mammary gland development and have an important role in lactation [126]. The Hedgehog signaling pathway is required for normal development of various mammalian organs. Although the research results on its role in mammary gland development have been inconsistent, the latest insights showed that it has an important role in mammary ductal morphogenesis [113]. Gene found in this category for FY included *PTCH1*, a known regulator of mammary ductal morphogenesis [127] that has never been associated with milk production traits in cattle thus far. *SCD*, a gene reported to participate in fatty acid synthesis in Italian Holstein and Simmental GWAS [128] was enriched in the fatty acid metabolism pathway as well for FY, which is in line with the aforementioned findings. Another gene enriched in the term “fatty acid metabolism” for FY was *HSD17B12*, previously reported as a candidate gene for fat yield [125]. The hippo signaling pathway regulates various biological processes in the organism, including potential role in pregnant and lactating mammary gland [114]. This pathway was enriched for the *NKD2*, gene described as a candidate gene for MY, FY, and PY in German Black Pied cattle [129]. Extracellular matrix (ECM) components are involved in mammary gland development processes as reviewed by Xu et al. [130]. Genes involved in this pathway, *THBS3* and *LAMA5*, were associated with MY. Both were previously reported as milk production and milk composition candidates [55, 131] and were found to be significantly enriched in the PI3K-Akt signaling pathway, as well. Many genes showed pleiotropic effects by involvement with the same terms across the three traits, which is in agreement with our candidate gene analysis, and previous research of other authors, as cited above.

The percentage of trait variance explained by the 50 most significant and 50 random variants from each chromosome, or so-called SNP-based heritability [132] was calculated to see how much of the genetic variance is attributable to top SNPs chosen for candidate gene research. To avoid overestimation of variance previously reported when using GREML [133], a GRM set up from 50K SNP chip data was included in the model, to account for further polygenic variance. Top SNPs explained much more variance than random ones (Table 2) indicating the potential presence of causal variants among those and underpinning the infinitesimal model. To eliminate multiple variants in high LD to each other that represent the same QTL, we pruned out the SNPs taking into account correlations between genotype allele counts [43]. Surprisingly, results differed depending on the trait; for MY, variants that were left after pruning explained more variance than the initial set of top SNPs. We expected that because the pruned variants spread over more QTL and should thus capture more variance. For FY and PY, however, pruned variants explained less variance than top SNPs. This could potentially be related to allelic heterogeneity in *DGAT1* because it can be assumed that the multiple variants capture more segregating variants [52].

## Conclusions

After performing large-scale GWAS we identified 30 new candidate genes for three milk production traits; MY (9), FY (8), and PY (18), of which 6 genes (*CDK5R2*, *STK25*, *PRDM1*, *KLHL12*, *RNF152*, and *FBXL19*) showed pleiotropic effects. These novel, functionally plausible candidates have not been reported for these traits so far. Variants located within or close to these genes explained a comparatively large proportion of genetic variance. In order to be able to fully exploit the power of GWAS, sequence data of very large samples are required, as shown in our study. Our findings add to existing knowledge of milk production traits architecture and convincingly demonstrate the power of our data set and strategy. Future studies incorporating health traits and their relationship with milk production may leverage the power of this data to add to the improvement of animal welfare.

## List of abbreviations

AF: allele frequency
apoB100: apolipoprotein B100
BTA: *Bos taurus* autosome
CNVR: copy number variation
DR2: dosage R-squared
DRPs: deregeressed proofs
FY: fat yield
GEBV: genomic estimated breeding values
GREML: genomic-relatedness-based restricted maximum-likelihood
GRM: genomic relationship matrix
GWAS: genome-wide association study
HD: high density
HPC: high-performance computing
KEGG: Kyoto Encyclopedia of Genes and Genomes
LD: linkage disequilibrium
MAF: minor allele frequency
MLMA: mixed linear model approach
MY: milk yield
N_e_: effective population size
NEFA: non-esterified fatty acids
ORA: over-representation analysis
PY: protein yield
SCC: somatic cell count
SCS: somatic cell score
VLDL: very low-density lipoprotein
WGS: whole-genome sequence

## Declarations

### Ethics approval and consent to participate

Not applicable. No live animals or animal material have been used in this study.

### Consent for publication

Not applicable.

### Availability of data and materials

The SNP chip genotype data and deregressed proofs are not available because they are the property of the national computing center in Germany (Vereinigte Informationssysteme Tierhaltung w.V.). Imputed genotypes and summary statistics will be provided upon reasonable request.

### Authors’ contributions

AMK performed the imputation, GWAS, and downstream analyses and wrote the paper. CR performed the genotype liftover and participated in genomic inflation analyses. JH provided the 50K SNP chip dataset, JP provided the HD reference dataset, and ZL provided the DRPs and gave useful comments. CFG participated in imputation and downstream analyses. CFG and JT supervised the study and participated in the writing of the paper. JT, JB, and GT conceived and supervised the project. All authors have read and approved the final manuscript.

## Acknowledgements

The authors want to thank Iona MacLeod from Agriculture Victoria Research, AgriBio, Centre for AgriBioscience, 5 Ring Road, LaTrobe University, Bundoora, Australia, and Donagh Berry from Teagasc, Animal & Grassland Research and Innovation Centre, Moorepark, Fermoy P61 P302, Co. Cork, Ireland for giving the approval for the use of the HD dataset. We also want to acknowledge the 1000 Bulls Genome Consortium for providing the Run9 WGS dataset.

## Additional files

**Additional file 1**

Format: .pdf

**Additional file 1 Table S1**

Title: Genotype arrays used for samples genotyping

**Additional file 1 Table S2**

Title: Composition of breeds of WGS reference panel

**Additional file 1 Table S3**

Title: Candidate genes associated with the top 50 variants for MY, FY, and PY

**Additional file 1 Table S4**

Title: Common genes between the three milk production traits

**Additional file 1 Table S5**

Title: Number of variant effects by type

**Additional file 1 Table S6**

Title: Functional Enrichment Analysis results for MY, FY, and PY

**Additional file 1 Table S7**

Title: List of known milk production and composition candidate genes identified in our study

**Additional file 1 Figure S1**

Format: .png

Title: Concordance between imputed KuhVision AF and 1000 Bulls Run9 AF from BTA16

**Additional file 2**

Format: .xlsx

**Additional file 2 Table S1**

Title: List of top variants for MY

**Additional file 2 Table S2**

Title: List of top variants for FY

**Additional file 2 Table S3**

Title: List of top variants for PY

**Additional file 3**

Format: .xlsx

**Additional file 3 Table S1**

Title: List of all genome-wide significant variants and their effects for MY

**Additional file 3 Table S2**

Title: List of all genome-wide significant variants and their effects for FY

**Additional file 3 Table S3**

Title: List of all genome-wide significant variants and their effects for PY

**Additional file 4**

Format: .xlsx

**Additional file 4 Table S1**

Title: Genes whose functions couldn’t be linked with milk production traits

