## Additional file 1 for "Sequence-based GWAS in 180 000 German Holstein cattle reveals new candidate genes for milk production traits"

**Table S1 Genotype arrays used for samples genotyping**

| Genotype array | SNP number on array | Number of animals genotyped |
|---|---|---|
| AffyVM2 | 63836 | 1 |
| AM3 | 61792 | 1 |
| AxiomMD V3 | 65004 | 487 |
| EuroG MD | 49331 | 4661 |
| EuroG MD V1.1 | 49852 | 2517 |
| EuroG MD V2 | 54271 | 1226 |
| EuroG MD V3 | 62902 | 3 |
| EuroG10K | 9072 | 440 |
| EuroG10K V1.1 | 9075 | 463 |
| EuroG10K V2 | 9001 | 614 |
| EuroG10K V4 | 11490 | 2856 |
| EuroG10K V5 | 13787 | 94423 |
| EuroG10K V7 | 13329 | 104594 |
| EuroG10K V8 | 13674 | 39156 |
| GGP3 | 26151 | 1 |
| GGP4 | 30105 | 27 |
| GMD | 47843 | 5 |
| Illumina 50k V2 | 54609 | 281 |
| Illumina 50k V3 | 53218 | 348 |
| Illumina LD | 6909 | 2 |
| WeatherbysVersa1.3 | 49629 | 95 |

**Table S2 Composition of breeds of WGS reference panel**

| Breed | Animal number |
|---|---|
| Abondance | 9 |
| Alentejana | 1 |
| Altai | 20 |
| ANAN | 2 |
| Angus | 401 |
| AngusGerman | 1 |
| AngusLowline | 4 |
| AngusRed | 33 |
| AngusSimmental | 1 |
| ARARCHAR | 1 |
| Aubrac | 9 |
| Auroch | 1 |
| AyrshireFinnish | 45 |
| BeefShorthorn | 8 |
| BelgianBlue | 9 |
| BelgianBlueHolstein | 4 |
| BelgianBlueLimousin | 1 |
| BelgianRedWhiteCampine | 10 |
| BelgiumBlue | 1 |
| BeltedCattle | 1 |
| BlondedAquitaine | 41 |
| Bohuskulla | 3 |
| Boskarin | 1 |
| Braunvieh | 6 |
| BrownSwiss | 294 |
| Buryat | 19 |
| Busa | 10 |
| Cabannina | 2 |
| Charolais | 154 |
| CharolaisAngus | 1 |
| CharolaisRedAngus | 1 |
| ChiAngus | 1 |
| Chianina | 15 |
| Cloned-polledDairyBull | 2 |
| Composite | 1 |
| Corriente | 4 |
| CostenoConCuernos | 2 |

| | |
|---|---|
| Crossbreed | 35 |
| Crossbreed(25%RedAngus;25%Simmental;25%Gelbvieh;25%Hereford) | 1 |
| Crossbreed(37.5%Gelbvieh;25%RedAngus;12.5%Simmental;12.5%Limousin) | 1 |
| Crossbreed(50%Simmental;25%RedAngus;25%Charolais) | 1 |
| Crossbreed(50%Simmental;37.5%RedAngus;12.5%Angus) | 1 |
| Crossbreed(50%Simmental;50%RedAngus) | 1 |
| Crossbreed(62.5%Angus;12.5%Simmental;12.5%Gelbvieh;12.5%Hereford) | 1 |
| Crossbreed(75%Gelbvieh;25%Limousin) | 1 |
| Crossbreed(HO62.5%;MO25%;JE12.5%) | 1 |
| DanishRedDairy | 4 |
| DanishRedDairyHolstein | 1 |
| DeepRedCattle | 9 |
| DeutschesSchwarzbuntesNiederungsrind | 56 |
| Devon | 1 |
| Dexter | 2 |
| DutchBelted | 11 |
| DutchFriesianRed | 11 |
| DutchImprovedRed | 9 |
| EasternBelgianRedWhite | 7 |
| EasternFinncattle | 15 |
| EasternFlandersWhiteRed | 12 |
| Eringer | 4 |
| Evolène | 3 |
| FinnishAyrshire | 12 |
| Fjäll | 11 |
| FjällCattle | 6 |
| Fleckvieh | 162 |
| FriesianJersey | 9 |
| Galloway | 1 |
| GallowayBelted | 3 |
| Gelbvieh | 52 |
| GermanRedAngler | 6 |
| GreyCattle | 6 |
| GroningenWhiteHeaded | 10 |
| Guernsey | 20 |
| Hanwoo | 22 |
| Hasake | 6 |
| HEAN | 5 |
| Hereford | 141 |
| HerefordMiniature | 2 |
| HerefordPolled | 4 |

| | |
|---|---|
| Hinterwaelder | 3 |
| Holstein | 1148 |
| HolsteinCharolais | 22 |
| HolsteinFriesian | 41 |
| HolsteinHereford | 9 |
| HolsteinLimousinF1Crossbred | 2 |
| HolsteinRed | 20 |
| HolsteinSimmental | 4 |
| HolsteinxJerseyF1Crossbred | 1 |
| IcelandicCattle | 5 |
| Illawarra | 1 |
| ImprovedRed | 1 |
| JapaneseNative | 8 |
| Jersey | 195 |
| JerseyHolstein | 42 |
| JerseyLimousin | 2 |
| Jutland | 5 |
| Kalmyk | 3 |
| Kalmykian | 10 |
| Kazakh | 9 |
| KazakhWhiteheaded | 5 |
| Kholmogory | 32 |
| LatvianBrown | 10 |
| Limia | 1 |
| Limonero | 6 |
| Limousin | 101 |
| LimousinBrownSwiss | 1 |
| LimousinHereford | 2 |
| LimousinHolstein | 1 |
| LimousinSimmental | 2 |
| LithuanianRed | 12 |
| Lowline | 1 |
| Luxi | 1 |
| MAARCTwinner | 4 |
| MaineAnjou | 22 |
| Marchigiana | 9 |
| Maremmana | 1 |
| Maronesa | 1 |
| Menggu | 11 |
| MeuseRhineYssel | 25 |
| ModernAngler | 20 |

| | |
|---|---|
| ModernDanishRed | 54 |
| Mongolian | 4 |
| Montbeliarde | 63 |
| MurrayGrey | 2 |
| MWFDEU | 3 |
| Normande | 44 |
| NorthernFinncattle | 19 |
| NorwegianRed | 347 |
| OriginalBraunvieh | 125 |
| Ottonese | 2 |
| Pajuna | 1 |
| Parthenaise | 2 |
| PezzataRossaItaliana | 1 |
| Piedmontese | 10 |
| PiedmonteseNormande | 1 |
| Pinzgauer | 1 |
| PodolianSerbia | 10 |
| Podolica | 1 |
| PolishRed | 7 |
| RedAngus | 2 |
| RedDairy | 2 |
| RedWhiteDualPurpose | 17 |
| Rendena | 2 |
| Ringamålako | 8 |
| Rödkulla | 9 |
| Romagnola | 25 |
| RotesHöhenvieh | 6 |
| RougeDesPres | 9 |
| Salers | 25 |
| SanMartinero | 2 |
| Sayaguesa | 1 |
| ScottishHighland | 7 |
| Shorthorn | 33 |
| Sikias | 1 |
| Simmental | 137 |
| SimmentalAngus | 1 |
| SimmentalFleckviehPezzatarossa | 36 |
| SMSMCHAR | 1 |
| Stabilizer | 2 |
| SwedishPolled | 6 |
| SwedishRed | 50 |

| | |
|---|---|
| SwedishRedPolled | 6 |
| SwissFleckvieh | 10 |
| Tarentaise | 12 |
| TexasLonghorn | 3 |
| TraditionalAngler | 5 |
| TraditionalDanishRed | 15 |
| TraditionalLithuanianRed | 4 |
| TuranoMongolicus | 1 |
| Tuxer | 1 |
| TyroleanGrauvieh | 9 |
| TyroleanGrey | 17 |
| UkrainianGrey | 8 |
| Unknown | 199 |
| Väneko | 11 |
| Vorderwälder | 13 |
| Vosgienne | 4 |
| Wagyu | 29 |
| WagyuModern | 1 |
| WesternFinncattle | 15 |
| WestVlaamsRood | 11 |
| Xizang | 2 |
| Yakut | 44 |
| Yanbian | 11 |
| Yaroslavl | 22 |

**Table S3 Candidate genes associated with the top 50 variants for MY, FY, and PY**

| Trait | Chr | Genes |
|---|---|---|
| | 1 | *SLC37A1*, *PDE9A* |
| | 2 | *ENSBTAG00000051006*, *TNS1*, *CDK5R2*, *FEV* |
| | 3 | *EFNA4*, *DPM3*, *EFNA1*, *EFNA3*, *DCST1*, *ZBTB7B*, *PKLR*, *FDPS*, *FLAD1*, *KRTCAP2*, *DPM3*, *MTX1*, *THBS3*, *HCN3*, *CLK2*, *ADAM15*, *TRIM46*, *MUC1*, *THBS3* |
| | 4 | *ENSBTAG00000051416*, U6, *CDK6* |
| | 5 | *MGST1*, *SLC15A5* |
| | 6 | *NPFFR2*, *GC*, *ENSBTAG00000049290*, *SLC4A4* |

| | | |
|---|---|---|
| **MY** | 7 | *CD70*, *TNFSF9*, *VAV1*, *ADGRE1*, *ENSBTAG00000052522*, *ENSBTAG00000055049*, *CLPP*, *ACER1* |
| | 8 | *MPDZ*, *LURAP1L*, *ENSBTAG00000044352*, 7SK, *ENSBTAG00000055211*, *ENSBTAG00000053688*, *ENSBTAG00000049421* |
| | 9 | *PRDM1*, *ENSBTAG00000054637* |
| | 10 | *ENSBTAG00000051430*, *NREP* |
| | 11 | *ENSBTAG00000048091*, *PAEP*, *GLT6D1*, *LCN9* |
| | 12 | *CYSLTR2*, *ENSBTAG00000048524*, *ENSBTAG00000050275*, *SUCLA2*, *ENSBTAG00000052225* |
| | 13 | *LAMA5*, *GATA5*, *RBBP8NL*, *RPS21*, *CABLES2*, *MKX* |
| | 14 | *ADCK5*, *CPSF1*, *SLC52A2*, *SLC39A4*, *FBXL6*, *TMEM249*, *SCRT1* |
| | 15 | *RAB6A*, U6, *MRPL48* |
| | 16 | *ENSBTAG00000052913*, *SOX13*, *PLEKHA6*, *CASZ1*, *NFASC*, *SYT2*, *ENSBTAG00000049657*, *ENSBTAG00000052560*, *ZNF281*, *PPP1R12B* |
| | 17 | *CRYBB2*, *GRK3* |
| | 19 | *PELP1*, *TUBG1*, *TUBG2*, bta-mir-2338, *CAVIN1*, *STAT5B*, *ZFP3*, *KIF1C*, *STAT3*, *KAT2A*, *DHX58*, *CXCL16*, *MED11*, *RAB5C*, *LPO*, *TM4SF5*, *ZMYND15*, *ENSBTAG00000053969*, *PLD2*, *PSMB6*, *HCRT*, *KCNH4*, *STAT5A*, *ZNF385C*, *CCR10*, *CNTNAP1* |
| | 20 | *GHR*, *OXCT1*, *FBXO4*, *C20H5orf51*, *ZNF131*, *ENSBTAG00000054352*, *ENSBTAG00000052195* |
| | 23 | *ENSBTAG00000055279*, *ENSBTAG00000055288*, *ENSBTAG00000054889*, bta-mir-7857-1, *SLC17A4*, *HIST1H2AG*, *ENSBTAG00000050532*, *ENSBTAG00000054340*, *ENSBTAG00000054480*, bta-mir-2379, *OR2B6*, *ENSBTAG00000039145*, *ZNF322*, *ENSBTAG00000054898*, *ENSBTAG00000039274* |
| | 24 | *RNF152*, *PIGN*, *ZNF521*, *HRH4*, *ENSBTAG00000054502*, *ENSBTAG00000047569* |
| | 25 | *FBXL19*, *ENSBTAG00000051451*, *SETD1A*, *PHKG2*, *TMEM265*, U6, *SPN*, *ENSBTAG00000046752*, bta-mir-2385, *PAGR1*, *KIF22*, *MAZ*, *C25H16orf54*, *QPRT*, *KCTD13*, *EIF3CL*, *CLN3*, *ENSBTAG00000043974*, *ALDOA*, bta-mir-12060, *ENSBTAG00000050743*, *MVP*, *ZNF688*, *DOC2A*, *INO80E* |
| | 27 | *GOLGA7*, *GINS4*, *GPAT4*, *NKX6-3*, *ENSBTAG00000027629*, *ENSBTAG00000054394*, *ENSBTAG00000003275*, *SFRP1*, 5S rRNA |

| | | |
|---|---|---|
| | 28 | *ENSBTAG00000048611*, *ENSBTAG00000053285*, *ZMIZ1*, *ZNF365*, *ENSBTAG00000051468* |
| | 29 | *KMT5B*, *C29H11orf24*, *IGF2*, *KCNQ1*, *ASCL2*, *TH* |
| **FY** | 1 | *IGF2BP2*, *TRA2B*, *SENP2*, *LIPH* |
| | 2 | *GMPPA*, *ASIC4*, *ENSBTAG00000052917*, *TNS1*, *ENSBTAG00000051006*, *NHEJ1*, *TUBA4A*, *TUBA1D*, *CFAP65*, *PAFAH2*, *SPEG*, *DNPEP*, *DES*, *STMN1*, *CYP27A1* |
| | 3 | *FCRLA*, *FCGR2B*, bta-mir-2285t, *FCRLB*, *DUSP12*, *STK25*, *BOK*, *HDLBP*, *GAL3ST2*, *NEU4*, *THAP4*, *SEPTIN2* |
| | 4 | *ENSBTAG00000054159*, *ENSBTAG00000052920* |
| | 5 | *MGST1*, *SLC15A5* |
| | 6 | *GC*, *NPFFR2*, *ENSBTAG00000049290* |
| | 8 | *PTCH1*, *ENSBTAG00000049821* |
| | 10 | *HOMER1*, *FLVCR2* |
| | 11 | *ENSBTAG00000044370*, *LPIN1* |
| | 12 | *CYSLTR2*, *ENSBTAG00000048524*, *LRCH1*, *ENSBTAG00000049836*, *ENSBTAG00000026070*, *ENSBTAG00000046041* |
| | 14 | *CPSF1*, *SLC39A4*, *ADCK5*, *TMEM249*, *SCRT1*, *SLC52A2*, *FBXL6*, *ENSBTAG00000053637* |
| | 15 | *EHF*, *APIP*, *HSD17B12*, *ALKBH3*, *KBTBD3*, *MSANTD4*, *ELF5*, *AASDHPPT*, *TTC17*, *GRIA4* |
| | 16 | *ENSBTAG00000052560*, *ZNF281*, *KLHL12*, *RABIF*, *ENSBTAG00000016189*, *PPP1R12B*, *ENSBTAG00000049594*, *ELF3*, *RNPEP*, *TIMM17A*, *PKP1*, *IGFN1*, *SYT2*, *LGR6*, *PTPN7*, *ENSBTAG00000055184* |
| | 17 | *MED13L*, *ENSBTAG00000052624* |
| | 18 | *ENSBTAG00000021433*, *ENSBTAG00000054418*, *ENSBTAG00000048680*, *ENSBTAG00000052289*, *HAS1*, *ENSBTAG00000039491*, bta-mir-125a, *SPACA6*, *MIR99B*, *MIRLET7E*, *ENSBTAG00000052124*, *ENSBTAG00000050359*, *ENSBTAG00000050920*, *ZNF613*, *ENSBTAG00000051367*, *ZNF614*, *ZNF432*, *ZNF350*, *ENSBTAG00000050488*, *ENSBTAG00000054038*, *DYNLRB2*, *ENSBTAG00000054728*, *ENSBTAG00000051227*, *ENSBTAG00000049640*, *ENSBTAG00000054322*, *ENSBTAG00000050562*, *ENSBTAG00000011844*, *ENSBTAG00000045880* |
| | 19 | *CCDC57* |

| | | |
|---|---|---|
| | 20 | *GHR, FBXO4, GABRP, RANBP17* |
| | 22 | *ENSBTAG00000052462, KLF15, CFAP100* |
| | 23 | *DDX39B, ENSBTAG00000031913, ENSBTAG00000013919, ENSBTAG00000048364, ENSBTAG00000026163, ENSBTAG00000005146, BOLA-NC1, MCCD1, JSP.1* |
| | 25 | *ENSBTAG00000031582, VKORC1L1, ENSBTAG00000054461, ITGAD, SLC5A2, TGFB1I1* |
| | 26 | *PAX2, SNORA70, SCD,* bta-mir-12016, *SLF2, HIF1AN* |
| | 27 | *ENSBTAG00000024530, THRB* |
| | 28 | *ENSBTAG00000051468, ENSBTAG00000053285, ZMIZ1* |
| | 29 | *KCNQ1, ASCL2, TH, IGF2, KIRREL3, ENSBTAG00000052359, ENSBTAG00000054745, CCDC15, SLC37A2, ENSBTAG00000053793, ENSBTAG00000019579, CARS1, NAP1L4, TMEM218, ENSBTAG00000048480, TOLLIP, ST3GAL4* |
| **PY** | 1 | *ENSBTAG00000053000, ENSBTAG00000046447, RUNX1, ENSBTAG00000030105, ENSBTAG00000054155, ENSBTAG00000052599* |
| | 2 | *ENSBTAG00000051006, TNS1, CDK5R2, FEV, TUBA4A, TUBA1D, TNP1, ENSBTAG00000048421, ENSBTAG00000048671, CYBRD1* |
| | 3 | *ENSBTAG00000053053, ENSBTAG00000013507, ENSBTAG00000050002, ENSBTAG00000052229, CRNN, ENSBTAG00000054258, ENSBTAG00000054333, ENSBTAG00000048997, FARP2, STK25, ENSBTAG00000048451, ENSBTAG00000053914, ENSBTAG00000039357, BOK, THAP4, ENSBTAG00000050431, CRCT1, C3H1orf68, RPTN* |
| | 4 | *ENSBTAG00000051416,* U6, *SUGCT* |
| | 5 | *ABCC9, ST8SIA1, ENSBTAG00000026611, CMAS* |
| | 6 | *GC, NPFFR2, ENSBTAG00000049290, SLC4A4* |
| | 7 | *ENSBTAG00000012150, EFNA2, PWWP3A, ENSBTAG00000050222, UHRF1, ADAMTSL5, ENSBTAG00000049103, ENSBTAG00000049174, MEX3D, MBD3, UQCR11, CD70, TNFSF9, MIDN, APC2, ARRDC5* |
| | 8 | *ENSBTAG00000044352,* 7SK |
| | 9 | *LIN28B, ENSBTAG00000054637, PRDM1, PREP, ENSBTAG00000048760, ATG5, POPDC3, BVES, CRYBG1, HACE1, ENSBTAG00000053858,* U6 |

| | | |
|---|---|---|
| | 10 | *ARSB*, *MAP2K5*, *ITGA11*, *CORO2B*, *RPLP1*, *ENSBTAG00000052439*, *YTHDC2*, *ENSBTAG00000049590*, *HOMER1* |
| | 11 | *PAEP*, *GLT6D1*, *ENSBTAG00000048091* |
| | 12 | *CYSLTR2*, *ENSBTAG00000048524*, *ENSBTAG00000055106*, *ENSBTAG00000053737*, *FNDC3A*, *ENSBTAG00000050275*, *SUCLA2*, *CAB39L*, *ENSBTAG00000052225*, *RB1* |
| | 13 | *BPIFB3*, *PARD3*, *GPCPD1*, *ENSBTAG00000054005* |
| | 14 | *ADCK5*, *CPSF1*, *FBXL6*, *SLC52A2*, *TMEM249*, *SLC39A4*, *SCRT1* |
| | 16 | *ENSBTAG00000052560*, *ZNF281*, *CAMSAP2*, *KLHL12*, *IGFN1*, *KDM5B*, *PPP1R12B*, *RABIF*, *ENSBTAG00000016189*, *PKP1*, *GPR25*, *INAVA*, *ENSBTAG00000049594*, *ELF3*, *RNPEP*, *TIMM17A*, *SYT2*, *ENSBTAG00000049657* |
| | 17 | *CRYBB2*, *GRK3*, *KSR2*, *NOS1*, *ENSBTAG00000031468*, *ENSBTAG00000053312*, *HORMAD2*, *LIF*, *RPH3A* |
| | 18 | *ZNF423*, *ENSBTAG00000051264*, *ENSBTAG00000051570*, *CBLN1*, *ENSBTAG00000050940*, *WWP2* |
| | 19 | *PLD2*, *PSMB6*, *PELP1*, bta-mir-195, bta-mir-497, *BCL6B*, *RNASEK*, C19H17orf49, *MINK1*, bta-mir-2338, *ZMYND15*, *CXCL16*, *MED11*, *TM4SF5*, *ZFP3*, *KIF1C*, *DERL2*, *DHX33*, *SPAG7*, *ENO3*, *ARRB2*, *ALOX15*, *ENSBTAG00000013906*, *ENSBTAG00000048823*, *CAMTA2*, *ENSBTAG00000055281*, *ENSBTAG00000050735*, *NLRP1*, *ENSBTAG00000053786*, *ALOX12* |
| | 20 | *ENSBTAG00000054687*, *ENSBTAG00000052982*, *SLC9A3*, 5S rRNA, *AHRR*, *EXOC3*, *ENSBTAG00000026527*, *NKD2*, *ENSBTAG00000049710*, *SLC12A7*, *DOCK2*, *FOXI1*, *CCDC127*, *SDHA*, *LRRC14B*, *PDCD6*, *GABRP*, *RANBP17*, *PTGER4*, U2, *ENSBTAG00000050512*, *IRX1*, *ENSBTAG00000052104*, *PIK3R1*, *NSG2*, bta-mir-584-6 |
| | 23 | *ENSBTAG00000006339*, *ENSBTAG00000027279*, *ENSBTAG00000055279*, *ENSBTAG00000055288*, *ZNF391*, *POM121L2*, *ENSBTAG00000000228*, *ENSBTAG00000037628*, *ENSBTAG00000050532*, *HIST1H2AG*, *ENSBTAG00000054340*, *ZNF184*, *ENSBTAG00000050455*, *ENSBTAG00000054760* |
| | 24 | *RNF152*, *PIGN*, *BCL2*, *ENSBTAG00000054502*, *ENSBTAG00000047569* |
| | 25 | *FBXL19*, *ENSBTAG00000054461*, *ITGAD*, *ENSBTAG00000031582*, *VKORC1L1*, *TGFB1I1*, *SLC5A2*, *ZNF629*, *BCL7C* |
| | 26 | *LIPK*, *LIPN*, *TACC2*, *C26H10orf120*, *ENSBTAG00000039755*, *RNLS*, *DHX32*, *SPADH2*, *SPADH1*, *KIF20B*, *ENSBTAG00000020790*, U7, *ENSBTAG00000048815*, |

| | | |
|---|---|---|
| | | *ATAD1*, *PTEN*, *ENSBTAG00000051599*, *ENSBTAG00000023846*, *BCCIP*, *EDRF1*, *ENSBTAG00000050262*, *ENSBTAG00000054811* |
| | 27 | *GINS4*, *GPAT4*, *GOLGA7*, *NKX6-3*, *ENSBTAG00000027629*, *ENSBTAG00000054394*, *ENSBTAG00000003275* |
| | 28 | *ENSBTAG00000051468*, *ENSBTAG00000053285*, *ZMIZ1* |
| | 29 | *KCNQ1*, *ENSBTAG00000048480*, *CCDC15*, *SLC37A2*, *KMT5B*, *TMEM218*, u6-2285r, *PKNOX2*, *EI24*, *STT3A*, *MSANTD2*, *ROBO3* |

**Table S4 Common genes between the three milk production traits**

| Traits | Genes | Number |
|---|---|---|
| MY, FY, PY | *NPFFR2*, *GC*, *SLC39A4*, *ZNF281*, *PPP1R12B*, *FBXL6*, *ENSBTAG00000049290*, *ENSBTAG00000051468*, *ZMIZ1*, *CPSF1*, *SYT2*, *SCRT1*, *ENSBTAG00000051006*, *CYSLTR2*, *TNS1*, *SLC52A2*, *ADCK5*, *ENSBTAG00000053285*, *ENSBTAG00000048524*, *ENSBTAG00000052560*, *KCNQ1*, *TMEM249* | 22 |
| MY, FY | *FBXO4*, *TH*, *SLC15A5*, *ASCL2*, *IGF2*, *GHR*, *MGST1* | 7 |
| MY, PY | *ENSBTAG00000054340*, *ENSBTAG0000005528*, *ENSBTAG00000055279*, *MED11*, *PELP1*, *ENSBTAG00000044352*, *PLD2*, *ZFP3*, *ENSBTAG00000052225*, *ENSBTAG00000054637*, 7SK, *FEV*, *ZMYND15*, *ENSBTAG00000054502*, *GPAT4*, *ENSBTAG00000054394*, *GINS4*, *KMT5B*, 5S rRNA, *NKX6-3*, *CDK5R2*, *TNFSF9*, *RNF152*, *ENSBTAG00000027629*, *PSMB6*, *CRYBB2*, U6, *GRK3*, *ENSBTAG00000048091*, *ENSBTAG00000051416*, *CXCL16*, *PIGN*, | 49 |

| | | |
|---|---|---|
| | *HIST1H2AG*, bta-mir-2338, *ENSBTAG00000003275*, *ENSBTAG00000050275*, *SLC4A4*, *ENSBTAG00000047569*, *TM4SF5*, *SUCLA2*, *GLT6D1*, *KIF1C*, *ENSBTAG00000050532*, *FBXL19*, *PAEP*, *CD70*, *PRDM1*, *ENSBTAG00000049657*, *GOLGA7* | |
| FY, PY | *TUBA1D*, *ENSBTAG00000031582*, *CCDC15*, *TUBA4A*, *THAP4*, *RABIF*, *TIMM17A*, *PKP1*, *VKORC1L1*, *ENSBTAG00000016189*, *STK25*, *ITGAD*, *SLC5A2*, *ENSBTAG00000048480*, *BOK*, *TGFB1I1*, *HOMER1*, *SLC37A2*, *IGFN1*, *ENSBTAG00000054461*, *ELF3*, *KLHL12*, *GABRP*, *RANBP17*, *RNPEP*, *ENSBTAG00000049594*, *TMEM218* | 27 |

### Table S5 Number of variant effects by type

| Predicted effect | MY | FY | PY |
|---|---|---|---|
| 3 prime UTR variant | 492/0.504% | 480/0.629% | 343/0.524% |
| 5 prime UTR premature start codon gain variant | 36/0.037% | 32/0.042% | 17/0.026% |
| 5 prime UTR variant | 256/0.262% | 212/0.278% | 164/0.251% |
| Conservative inframe deletion | 3/0.003% | 0 | 0 |
| Conservative inframe insertion | 1/0.001% | 4/0.005% | 3/0.005% |
| Disruptive inframe deletion | 7/0.007% | 8/0.01% | 8/0.012% |

| | | | |
|---|---|---|---|
| Disruptive inframe insertion | 5/0.005% | 5/0.007% | 0 |
| Downstream gene variant | 6462/6.613% | 4789/6.274% | 4454/6.806% |
| Frameshift variant | 5/0.005% | 11/0.014% | 5/0.008% |
| Intergenic region | 35,983/36.826% | 28,028/36.721% | 25,419/38.843% |
| Intron variant | 45,883/46.958% | 35,924/47.066% | 29,585/45.209% |
| Missense variant | 446/0.456% | 442/0.579% | 293/0.448% |
| Noncoding transcript exon variant | 98/0.1% | 72/0.094% | 33/0.05% |
| Splice acceptor variant | 1/0.001% | 2/0.003% | 2/0.003% |
| Splice donor variant | 2/0.002% | 4/0.005% | 3/0.005% |
| Splice region variant | 136/0.139% | 108/0.141% | 89/0.136% |
| Start lost | 1/0.001% | 2/0.003% | 1/0.002% |
| Stop gained | 1/0.001% | 1/0.001% | 2/0.003% |
| Stop lost | 1/0.001% | 0 | 0 |
| Stop retained variant | 1/0.001% | 0 | 0 |
| Synonymous variant | 866/0.886% | 750/0.983% | 634/0.969% |
| Upstream gene variant | 7024/7.189% | 5453/7.144% | 4385/6.701% |

### Table S6 Functional Enrichment Analysis results for MY, FY, and PY

| Trait | ID | Description | Gene ratio | *p*-value | Gene ID |
|---|---|---|---|---|---|
| MY | bta00053 | Ascorbate and aldarate metabolism | 12/290 | 1.28787e-11 | *UGT2B10*, *MGC152010*, *UGT2A1*, *LOC615303*, *LOC100140261*, *LOC100138908*, *LOC540615*, *LOC100296421*, *LOC540544*, *LOC530553*, *LOC526979*, *LOC781988* |
| MY | bta00040 | Pentose and glucuronate interconversions | 12/290 | 1.30630e-10 | *UGT2B10*, *MGC152010*, *UGT2A1*, *LOC615303*, *LOC100140261*, *LOC100138908*, *LOC540615*, *LOC100296421*, *LOC540544*, *LOC530553*, *LOC526979*, *LOC781988* |
| MY | bta00140 | Steroid hormone biosynthesis | 16/290 | 1.27117e-09 | *UGT2B10*, *MGC152010*, *UGT2A1*, *SULT1E1*, *CYP11B1*, *HSD17B1*, *LOC615303*, *LOC100140261*, *LOC100138908*, *LOC540615*, *LOC100296421*, *LOC540544*, *LOC530553*, *LOC526979*, *LOC781988*, *LOC112449566* |
| MY | bta00860 | Porphyrin metabolism | 12/290 | 2.32597e-09 | *UGT2B10*, *MGC152010*, *UGT2A1*, *LOC615303*, *LOC100140261*, *LOC100138908*, *LOC540615*, *LOC100296421*, *LOC540544*, *LOC530553*, *LOC526979*, *LOC781988* |
| MY | bta00982 | Drug metabolism - cytochrome P450 | 13/290 | 4.16819e-08 | *MGST1*, *UGT2B10*, *MGC152010*, *UGT2A1*, *LOC615303*, *LOC100140261*, *LOC100138908*, *LOC540615*, *LOC100296421*, *LOC540544*, *LOC530553*, *LOC526979*, *LOC781988* |
| MY | bta00983 | Drug metabolism - other enzymes | 14/290 | 7.11112e-08 | *MGST1*, *UGT2B10*, *MGC152010*, *UGT2A1*, *RRM2B*, *LOC615303*, *LOC100140261*, *LOC100138908*, *LOC540615*, *LOC100296421*, *LOC540544*, *LOC530553*, *LOC526979*, *LOC781988* |
| MY | bta05204 | Chemical carcinogenesis - DNA adducts | 13/290 | 7.57564e-08 | *MGST1*, *UGT2B10*, *MGC152010*, *UGT2A1*, *LOC615303*, *LOC100140261*, *LOC100138908*, *LOC540615*, *LOC100296421*, *LOC540544*, *LOC530553*, *LOC526979*, *LOC781988* |
| MY | bta00980 | Metabolism of xenobiotics by cytochrome P450 | 13/290 | 1.10686e-07 | *MGST1*, *UGT2B10*, *MGC152010*, *UGT2A1*, *LOC615303*, *LOC100140261*, *LOC100138908*, *LOC540615*, *LOC100296421*, *LOC540544*, *LOC530553*, *LOC526979*, *LOC781988* |

| | | | | | |
|---|---|---|---|---|---|
| MY | bta00830 | Retinol metabolism | 13/290 | 2.69008e-07 | *UGT2B10*, *MGC152010*, *UGT2A1*, *DGAT1*, *LOC615303*, *LOC100140261*, *LOC100138908*, *LOC540615*, *LOC100296421*, *LOC540544*, *LOC530553*, *LOC526979*, *LOC781988* |
| MY | bta04976 | Bile secretion | 15/290 | 8.80114e-07 | *SLCO1B3*, *ABCG2*, *UGT2B10*, *MGC152010*, *UGT2A1*, *SLC4A4*, *LOC615303*, *LOC100140261*, *LOC100138908*, *LOC540615*, *LOC100296421*, *LOC540544*, *LOC530553*, *LOC526979*, *LOC781988* |
| MY | bta05207 | Chemical carcinogenesis - receptor activation | 22/290 | 9.25980e-07 | *CHRNB2*, *VDR*, *MGST1*, *UGT2B10*, *MGC152010*, *UGT2A1*, *MYC*, *ARRB2*, *STAT5B*, *STAT5A*, *STAT3*, *FGF10*, *RPS6KB2*, *LOC615303*, *LOC100140261*, *LOC100138908*, *LOC540615*, *LOC100296421*, *LOC540544*, *LOC530553*, *LOC526979*, *LOC781988* |
| MY | bta01240 | Biosynthesis of cofactors | 17/290 | 4.68627e-06 | *FLAD1*, *BCAT1*, *UGT2B10*, *MGC152010*, *UGT2A1*, *MTHFD2L*, *NAPRT*, *COASY*, *LOC615303*, *LOC100140261*, *LOC100138908*, *LOC540615*, *LOC100296421*, *LOC540544*, *LOC530553*, *LOC526979*, *LOC781988* |
| MY | bta04657 | IL-17 signaling pathway | 12/290 | 3.21767e-05 | *S100A7*, *S100A8*, *S100A9*, *CXCL8*, *CXCL5*, *CXCL2*, *CXCL3*, *GRO1*, *MAPK15*, *SRSF1*, *LOC786350*, *LOC112445860* |
| MY | bta04151 | PI3K-Akt signaling pathway | 26/290 | 0.00013 | *THBS3*, *EFNA1*, *EFNA3*, *EFNA4*, *IL6R*, *COL2A1*, *GYS2*, *SPP1*, *IBSP*, *EREG*, *AREG*, *TSC1*, *LAMA5*, *PTK2*, *MYC*, *YWHAZ*, *BRCA1*, *FGF10*, *GHR*, *PRKAA1*, *OSMR*, *RPS6KB2*, *IGF2*, *INS*, *NTRK1*, *PRLR* |
| MY | bta04061 | Viral protein interaction with cytokine and cytokine receptor | 9/290 | 0.00273 | *IL6R*, *CXCL8*, *CXCL5*, *CXCL2*, *CXCL3*, *GRO1*, *CCR10*, *CCL28*, *PPBP* |
| MY | bta04012 | ErbB signaling pathway | 8/290 | 0.004496 | *EREG*, *AREG*, *BTC*, *PTK2*, *MYC*, *STAT5B*, *STAT5A*, *RPS6KB2* |
| MY | bta04917 | Prolactin signaling pathway | 8/290 | 0.004496 | *SOCS2*, *CSN2*, *STAT5B*, *STAT5A*, *STAT3*, *TH*, *INS*, *PRLR* |
| MY | bta04062 | Chemokine signaling pathway | 13/290 | 0.00666 | *CXCL8*, *CXCL5*, *CXCL2*, *CXCL3*, *GRO1*, *PTK2*, *CXCL16*, *ARRB2*, *STAT5B*, *STAT3*, *CCR10*, *CCL28*, *PPBP* |
| MY | bta01230 | Biosynthesis of amino acids | 7/290 | 0.00742 | *PKLR*, *PFKM*, *BCAT1*, *GPT*, *PYCR3*, *ENO3*, *TALDO1* |
| MY | bta04950 | Maturity onset diabetes of the young | 4/290 | 0.00801 | *PKLR*, *IAPP*, *MAFA*, *INS* |

| | | | | | |
|---|---|---|---|---|---|
| MY | bta05134 | Legionellosis | 6/290 | 0.008994 | *CXCL8*, *CXCL2*, *CXCL3*, *GRO1*, *RAB1A*, *HSF1* |
| MY | bta04060 | Cytokine-cytokine receptor interaction | 18/290 | 0.01702 | *IL6R*, *CXCL8*, *CXCL5*, *CXCL2*, *CXCL3*, *GRO1*, *CD70*, *GDF6*, *CXCL16*, *CCR10*, *CCL28*, *GHR*, *OSMR*, *LIFR*, *CLCF1*, *PPBP*, *TNFSF9*, *PRLR* |
| MY | bta04512 | ECM-receptor interaction | 7/290 | 0.02185 | *THBS3*, *COL2A1*, *SPP1*, *IBSP*, *LAMA5*, *GP1BA*, *MEPE* |
| FY | bta00040 | Pentose and glucuronate interconversions | 12/234 | 1.06005e-11 | *UGT2B10*, *UGT2A1*, *DCXR*, *LOC615303*, *LOC100140261*, *LOC100138908*, *LOC540615*, *LOC100296421*, *LOC540544*, *LOC526979*, *LOC781988*, *LOC112441469* |
| FY | bta00053 | Ascorbate and aldarate metabolism | 10/234 | 7.28625e-10 | *UGT2B10*, *UGT2A1*, *LOC615303*, *LOC100140261*, *LOC100138908*, *LOC540615*, *LOC100296421*, *LOC540544*, *LOC526979*, *LOC781988* |
| FY | bta00860 | Porphyrin metabolism | 11/234 | 3.2303e-09 | *UGT2B10*, *UGT2A1*, *COX15*, *LOC615303*, *LOC100140261*, *LOC100138908*, *LOC540615*, *LOC100296421*, *LOC540544*, *LOC526979*, *LOC781988* |
| FY | bta00140 | Steroid hormone biosynthesis | 14/234 | 5.56522e-09 | *UGT2B10*, *UGT2A1*, *SULT1E1*, *CYP11B1*, *HSD17B12*, *LOC615303*, *LOC100140261*, *LOC100138908*, *LOC540615*, *LOC100296421*, *LOC540544*, *LOC526979*, *LOC781988*, *LOC112449566* |
| FY | bta00980 | Metabolism of xenobiotics by cytochrome P450 | 12/234 | 8.4126e-08 | *MGST1*, *UGT2B10*, *UGT2A1*, *DCXR*, *LOC615303*, *LOC100140261*, *LOC100138908*, *LOC540615*, *LOC100296421*, *LOC540544*, *LOC526979*, *LOC781988* |
| FY | bta00982 | Drug metabolism - cytochrome P450 | 11/234 | 3.26363e-07 | *MGST1*, *UGT2B10*, *UGT2A1*, *LOC615303*, *LOC100140261*, *LOC100138908*, *LOC540615*, *LOC100296421*, *LOC540544*, *LOC526979*, *LOC781988* |
| FY | bta04976 | Bile secretion | 14/234 | 3.66835e-07 | *SLCO1A2*, *SLCO1B3*, *UGT2B10*, *UGT2A1*, *SLC4A4*, *ABCC2*, *LOC615303*, *LOC100140261*, *LOC100138908*, *LOC540615*, *LOC100296421*, *LOC540544*, *LOC526979*, *LOC781988* |
| FY | bta05204 | Chemical carcinogenesis - DNA adducts | 11/234 | 5.37812e-07 | *MGST1*, *UGT2B10*, *UGT2A1*, *LOC615303*, *LOC100140261*, *LOC100138908*, *LOC540615*, *LOC100296421*, *LOC540544*, *LOC526979*, *LOC781988* |
| FY | bta01240 | Biosynthesis of cofactors | 16/234 | 1.23730e-06 | *GMPPA*, *BCAT1*, *UGT2B10*, *UGT2A1*, *MTHFD2L*, *NAPRT*, |

| | | | | | |
|---|---|---|---|---|---|
| | | | | | *COX15*, *LOC615303*, *LOC100140261*, *LOC100138908*, *LOC540615*, *LOC100296421*, *LOC540544*, *LOC526979*, *LOC781988*, *VKORC1L1* |
| FY | bta00830 | Retinol metabolism | 11/234 | 1.5555e-06 | *UGT2B10*, *UGT2A1*, *DGAT1*, *LOC615303*, *LOC100140261*, *LOC100138908*, *LOC540615*, *LOC100296421*, *LOC540544*, *LOC526979*, *LOC781988* |
| FY | bta00983 | Drug metabolism - other enzymes | 11/234 | 2.69816e-06 | *MGST1*, *UGT2B10*, *UGT2A1*, *LOC615303*, *LOC100140261*, *LOC100138908*, *LOC540615*, *LOC100296421*, *LOC540544*, *LOC526979*, *LOC781988* |
| FY | bta04742 | Taste transduction | 9/234 | 0.00022 | *TAS2R42*, *TAS2R46*, *T2R65A*, *T2R12*, *T2R10C*, *TAS2R7*, *PKD2L1*, *LOC782957*, *TAS2R8* |
| FY | bta05207 | Chemical carcinogenesis - receptor activation | 15/234 | 0.00035 | *MGST1*, *UGT2B10*, *UGT2A1*, *CACNA1S*, *MIRLET7E*, *ARRB2*, *LOC615303*, *LOC100140261*, *LOC100138908*, *LOC540615*, *LOC100296421*, *LOC540544*, *LOC526979*, *LOC781988*, *KRAS* |
| FY | bta04936 | Alcoholic liver disease | 11/234 | 0.00170 | *TRA2B*, *CXCL8*, *CXCL2*, *CXCL3*, *GRO1*, *ADIPOR1*, *ACADVL*, *FASN*, *FAS*, *SCD*, *LPIN1* |
| FY | bta04340 | Hedgehog signaling pathway | 6/234 | 0.00268 | *PTCH1*, *ARRB2*, *CSNK1D*, *BTRC*, *SUFU*, *IHH* |
| FY | bta04062 | Chemokine signaling pathway | 11/234 | 0.00871 | *CXCL8*, *CXCL5*, *CXCL2*, *CXCL3*, *GRO1*, *PTK2*, *CXCL16*, *ARRB2*, *RAC3*, *PPBP*, *KRAS* |
| FY | bta04814 | Motor proteins | 11/234 | 0.01291 | *TUBA4A*, *KIF1A*, *CAPZA3*, *KIFC2*, *KIF14*, *TNNT2*, *DYNLRB2*, *ACTA2*, *KIF20B*, *TUBA1D*, *KIF1C* |
| FY | bta01212 | Fatty acid metabolism | 5/234 | 0.0149 | *HSD17B12*, *ACADVL*, *FASN*, *SCD*, *ELOVL3* |
| FY | bta05134 | Legionellosis | 5/234 | 0.0149 | *CXCL8*, *CXCL2*, *CXCL3*, *GRO1*, *HSF1* |
| FY | bta04650 | Natural killer cell mediated cytotoxicity | 9/234 | 0.01805 | *KLRD1*, *KLRK1*, *RAC3*, *FAS*, *LOC100140174*, *LOC618565*, *FCGR3A*, *KRAS*, *KLRC1* |
| FY | bta04012 | ErbB signaling pathway | 6/234 | 0.01881 | *CDKN1B*, *EREG*, *AREG*, *PTK2*, *KRAS*, *BTC* |
| PY | bta00053 | Ascorbate and aldarate metabolism | 13/184 | 1.31544e-15 | *UGT2B10*, *MGC152010*, *UGT2A1*, *LOC100138004*, *LOC615303*, *LOC100138908*, *LOC100140261*, *LOC540615*, *LOC100296421*, *LOC540544*, *LOC530553*, *LOC526979*, *LOC781988* |

| | | | | | |
|---|---|---|---|---|---|
| PY | bta00040 | Pentose and glucuronate interconversions | 13/184 | 1.84757e-14 | *UGT2B10*, *MGC152010*, *UGT2A1*, *LOC100138004*, *LOC615303*, *LOC100138908*, *LOC100140261*, *LOC540615*, *LOC100296421*, *LOC540544*, *LOC530553*, *LOC526979*, *LOC781988* |
| PY | bta00860 | Porphyrin metabolism | 13/184 | 4.94883e-13 | *UGT2B10*, *MGC152010*, *UGT2A1*, *LOC100138004*, *LOC615303*, *LOC100138908*, *LOC100140261*, *LOC540615*, *LOC100296421*, *LOC540544*, *LOC530553*, *LOC526979*, *LOC781988* |
| PY | bta00140 | Steroid hormone biosynthesis | 16/184 | 1.3318e-12 | *UGT2B10*, *MGC152010*, *UGT2A1*, *SULT1E1*, *CYP11B1*, *LOC100138004*, *LOC615303*, *LOC100138908*, *LOC100140261*, *LOC540615*, *LOC100296421*, *LOC540544*, *LOC530553*, *LOC526979*, *LOC781988*, *LOC112449566* |
| PY | bta04976 | Bile secretion | 18/184 | 1.93111e-12 | *SLCO1A2*, *SLCO1B3*, *UGT2B10*, *MGC152010*, *UGT2A1*, *SLC4A4*, *SLC9A3*, *LOC100138004*, *LOC615303*, *LOC100138908*, *LOC100140261*, *LOC540615*, *LOC100296421*, *LOC540544*, *LOC530553*, *LOC526979*, *LOC781988*, *ATP1B2* |
| PY | bta00982 | Drug metabolism - cytochrome P450 | 14/184 | 1.06414e-11 | *MGST1*, *UGT2B10*, *MGC152010*, *UGT2A1*, *LOC100138004*, *LOC615303*, *LOC100138908*, *LOC100140261*, *LOC540615*, *LOC100296421*, *LOC540544*, *LOC530553*, *LOC526979*, *LOC781988* |
| PY | bta05204 | Chemical carcinogenesis - DNA adducts | 14/184 | 2.11963e-11 | *MGST1*, *UGT2B10*, *MGC152010*, *UGT2A1*, *LOC100138004*, *LOC615303*, *LOC100138908*, *LOC100140261*, *LOC540615*, *LOC100296421*, *LOC540544*, *LOC530553*, *LOC526979*, *LOC781988* |
| PY | bta00980 | Metabolism of xenobiotics by cytochrome P450 | 14/184 | 3.28581e-11 | *MGST1*, *UGT2B10*, *MGC152010*, *UGT2A1*, *LOC100138004*, *LOC615303*, *LOC100138908*, *LOC100140261*, *LOC540615*, *LOC100296421*, *LOC540544*, *LOC530553*, *LOC526979*, *LOC781988* |
| PY | bta00830 | Retinol metabolism | 14/184 | 9.20545e-11 | *UGT2B10*, *MGC152010*, *UGT2A1*, *DGAT1*, *LOC100138004*, *LOC615303*, *LOC100138908*, *LOC100140261*, *LOC540615*, *LOC100296421*, *LOC540544*, |

| | | | | | |
|---|---|---|---|---|---|
| | | | | | *LOC530553*, *LOC526979*, *LOC781988* |
| PY | bta00983 | Drug metabolism - other enzymes | 14/184 | 1.9762e-10 | *MGST1*, *UGT2B10*, *MGC152010*, *UGT2A1*, *LOC100138004*, *LOC615303*, *LOC100138908*, *LOC100140261*, *LOC540615*, *LOC100296421*, *LOC540544*, *LOC530553*, *LOC526979*, *LOC781988* |
| PY | bta01240 | Biosynthesis of cofactors | 17/184 | 6.7795e-09 | *BCAT1*, *UGT2B10*, *MGC152010*, *UGT2A1*, *MTHFD2L*, *NAPRT*, *VKORC1L1*, *LOC100138004*, *LOC615303*, *LOC100138908*, *LOC100140261*, *LOC540615*, *LOC100296421*, *LOC540544*, *LOC530553*, *LOC526979*, *LOC781988* |
| PY | bta05207 | Chemical carcinogenesis - receptor activation | 16/184 | 5.40187e-06 | *MGST1*, *UGT2B10*, *MGC152010*, *UGT2A1*, *RB1*, *ARRB2*, *LOC100138004*, *LOC615303*, *LOC100138908*, *LOC100140261*, *LOC540615*, *LOC100296421*, *LOC540544*, *LOC530553*, *LOC526979*, *LOC781988* |
| PY | bta04062 | Chemokine signaling pathway | 11/184 | 0.0014 | *CXCL8*, *CXCL5*, *CXCL2*, *CXCL3*, *GRO1*, *PTK2*, *CXCL16*, *ARRB2*, *PIK3R5*, *PIK3R6*, *PPBP* |
| PY | bta05134 | Legionellosis | 5/184 | 0.00556 | *CXCL8*, *CXCL2*, *CXCL3*, *GRO1*, *HSF1* |
| PY | bta04964 | Proximal tubule bicarbonate reclamation | 3/184 | 0.00885 | *SLC4A4*, *SLC9A3*, *ATP1B2* |
| PY | bta04657 | IL-17 signaling pathway | 6/184 | 0.01061 | *CXCL8*, *XCL5*, *CXCL2*, *CXCL3*, *GRO1*, *MAPK15* |
| PY | bta04061 | Viral protein interaction with cytokine and cytokine receptor | 6/184 | 0.01115 | *CXCL8*, *CXCL5*, *CXCL2*, *CXCL3*, *GRO1*, *PPBP* |
| PY | bta04390 | Hippo signaling pathway | 8/184 | 0.01208 | *MOB1B*, *AFP*, *RASSF6*, *AREG*, *SCRIB*, *DLG4*, *DVL2*, *NKD2* |

**Table S7 List of known milk production and composition candidate genes identified in the study**

| CHR | GENE | TRAIT | PREVIOUSLY REPORTED BY |
|---|---|---|---|
| 1 | *SLC37A1* | MY | MY, FP, PP [1, 2] |
| 1 | *PDE9A* | MY | PY, MY, FY [3, 4] |

| | | | |
|---|---|---|---|
| 1 | *IGF2BP2* | FY | MY [5] |
| 1 | *TRA2B* | FY | MY [6] |
| 1 | *RUNX1* | PY | colostrum albumin concentration [7] |
| 1 | *ENSBTAG00000046447* | PY | milk production traits [8] |
| 2 | *TNS1* | MY, FY, PY | PUFA [9] |
| 2 | *CDK5R2* | MY, PY | MY [10] |
| 2 | *FEV* | MY, PY | MY [10] |
| 2 | *GMPPA* | FY | milk fat globule size regulation [11] |
| 2 | *PAFAH2* | FY | LP [12] |
| 2 | *STMN1* | FY | milk production [13] |
| 2 | *CYP27A1* | FY | MY, PY, FP, PP [14] |
| 2 | *CYBRD1* | PY | differential MY [15] |
| 3 | *EFNA4* | MY | PP, LPC, PC, milk calcium content [16–19] |
| 3 | *DPM3* | MY | PP, LPC, milk calcium content [1, 17, 19] |
| 3 | *EFNA1* | MY | MY, FP, PP [20] |
| 3 | *EFNA3* | MY | PP, LPC, milk calcium content [16, 17, 19] |
| 3 | *DCST1* | MY | PP, LPC, milk calcium content [1, 17, 19] |
| 3 | *ZBTB7B* | MY | PP, LPC, milk calcium content [1, 17, 19] |
| 3 | *PKLR* | MY | MY, FY, PY, PP [21, 22] |
| 3 | *FDPS* | MY | milk calcium content, FP [19, 23] |
| 3 | *FLAD1* | MY | LPC, milk calcium content [17, 19] |
| 3 | *KRTCAP2* | MY | milk calcium content, LPC, PP [19, 24] |
| 3 | *MTX1* | MY | PP, milk calcium content, LPC [1, 19, 24] |
| 3 | *THBS3* | MY | PP, milk calcium content [1, 19] |
| 3 | *HCN3* | MY | PP, milk calcium content [1, 19] |
| 3 | *CLK2* | MY | PP, milk calcium content [1, 19] |
| 3 | *ADAM15* | MY | PP, LPC, PC, milk calcium content [1, 17–19] |
| 3 | *TRIM46* | MY | PP, LPC, milk calcium content [1, 17, 19] |
| 3 | *MUC1* | MY | PP, LPC, milk calcium content, MY, PY [1, 17, 19, 25] |
| 3 | *FCRLA* | FY | milk composition [24] |
| 3 | *FCGR2B* | FY | MY, PY, PP [26, 27] |
| 3 | bta-mir-2285t | FY | MY [28] |
| 3 | *HDLBP* | FY | lipoprotein metabolism [29] |
| 3 | *GAL3ST2* | FY | upregulated in mammary fat pad in calves fed with different nutrient intake level [30] |
| 4 | U6 | MY, PY | MY, FY, PY, FP, PP [16] |
| 4 | *CDK6* | MY | serum albumin concentration [7] |
| 5 | *MGST1* | MY, FY | MY, FY, PY, FC, PP, PC, LPC, LY [1, 18, 31] |
| 5 | *SLC15A5* | MY, FY | MY, FY, FP, PP [16, 20] |
| 5 | *ABCC9* | PY | FY, FP, MY, PY [20, 32, 33] |
| 5 | *ST8SIA1* | PY | FY, FP [20, 23, 33] |

| | | | |
|---|---|---|---|
| 6 | *GC* | MY, FY, PY | MY, FY, PY [20, 34] |
| 6 | *NPFFR2* | MY, FY, PY | MY, PY [20, 35] |
| 6 | *SLC4A4* | MY, PY | MY, FY, PY, PP [20] |
| 7 | *CD70* | MY, PY | PP [36] |
| 7 | *TNFSF9* | MY, PY | PP [36] |
| 7 | *VAV1* | MY | PP [36] |
| 8 | *LURAP1L* | MY | FY, FP [37] |
| 8 | 7SK | MY, PY | PY, FP [16] |
| 9 | *ATG5* | PY | autophagosome formation-related gene in cow's mammary gland [38] |
| 9 | U6 | PY | casein content [39] |
| 10 | *NREP* | MY | LPC [40] |
| 10 | *FLVCR2* | FY | FY [16] |
| 10 | *CORO2B* | PY | PP [41] |
| 10 | *ITGA11* | PY | milk production [42] |
| 10 | *RPLP1* | PY | MY [43] |
| 10 | *YTHDC2* | PY | milk minerals [44] |
| 11 | *PAEP* | MY, PY | FP, PP, FC [1, 2, 18] |
| 11 | *GLT6D1* | MY, PY | FP, milk oligosaccharides [1, 45] |
| 11 | *LCN9* | MY | FY [16] |
| 11 | *LPIN1* | FY | MY, FY, PY, FP, PP [46] |
| 11 | *ENSBTAG00000048091* | MY, PY | heat stress response of UFA [47] |
| 12 | *ENSBTAG00000026070* | FY | milk production [8] |
| 12 | *FNDC3A* | PY | MY, FY, PY [48] |
| 13 | *LAMA5* | MY | MY, PP, LP [20, 49] |
| 13 | *RBBP8NL* | MY | MY, PP [20] |
| 13 | *MKX* | MY | MY, PY [8] |
| 13 | *BPIFB3* | PY | PY [16] |
| 13 | *GPCPD1* | PY | FP [16] |
| 13 | *ENSBTAG00000054005* | PY | FP [16] |
| 14 | *ADCK5* | MY, FY, PY | FP, MY, FY, PP [9, 16, 50–52] |
| 14 | *CPSF1* | MY, FY, PY | MY, FY, FP, PP [53] |
| 14 | *SLC52A2* | MY, FY, PY | MY, FY, PY, FP, PP, LY [8, 24, 40] |
| 14 | *SLC39A4* | MY, FY, PY | MY, FY, PY, FP, PP, PC [8, 16, 18] |
| 14 | *FBXL6* | MY, FY, PY | MY, FY, PY [54, 55] |
| 14 | *TMEM249* | MY, FY, PY | MY, FY, PY [8, 54] |
| 14 | *SCRT1* | MY, FY, PY | MY, FY, PY, FC [18, 54] |
| 14 | *ENSBTAG00000053637* | FY | MY, FY, FP, PP [16] |
| 15 | *RAB6A* | MY | MY, milk composition [56, 57] |
| 15 | U6 | MY | MY [16] |
| 15 | *MRPL48* | MY | FY [53] |
| 15 | *EHF* | FY | FY [16, 20] |

| | | | |
|---|---|---|---|
| 15 | *HSD17B12* | FY | FY [16] |
| 15 | *ELF5* | FY | FY [20] |
| 15 | *AASDHPPT* | FY | FY, FP [20] |
| 15 | *GRIA4* | FY | FY, FP [20] |
| 16 | *SOX13* | MY | PP [58] |
| 16 | *ZNF281* | MY, FY, PY | fatty acid profile [59] |
| 16 | *PLEKHA6* | MY | MY [37] |
| 16 | *NFASC* | MY | IGF-1 concentration in serum [60] |
| 16 | *ELF3* | FY, PY | milk production [61] |
| 17 | *CRYBB2* | MY, PY | FP [16] |
| 17 | *KSR2* | PY | FY, PY, FP, PP [14] |
| 17 | *NOS1* | PY | MY, FY, PY [48] |
| 17 | *RPH3A* | PY | FP [62] |
| 18 | *ENSBTAG00000039491* | FY | FY, FP [37] |
| 18 | bta-mir-125a | FY | milk fat synthesis [63] |
| 18 | *ZNF613* | FY | fatty acid [59] |
| 18 | *ZNF432* | FY | colostrum quantity [64] |
| 18 | *ZNF614* | FY | colostrum quantity [64] |
| 18 | *ZNF350* | FY | fatty acid, colostrum quantity [59, 64] |
| 19 | *CCDC57* | FY | FY, FP, FC [20, 65] |
| 18 | *ZNF423* | PY | LY [17] |
| 19 | *PELP1* | MY, PY | MY, FY, PY [8] |
| 19 | *TUBG2* | MY | differentially expressed in mammary tissue of Holstein cows at early lactation and non-lactation [66] |
| 19 | bta-mir-2338 | MY, PY | MY, FY, PY [8] |
| 19 | *STAT5B* | MY | PP, MY, PY [16, 67] |
| 19 | *ZFP3* | MY, PY | FY, PY [8] |
| 19 | *KIF1C* | MY, PY | FY, PY [8] |
| 19 | *STAT3* | MY | PP [16] |
| 19 | *KAT2A* | MY | LP, PP, LC [24, 40] |
| 19 | *DHX58* | MY | PP, LC [1, 40] |
| 19 | *CXCL16* | MY, PY | MY, FY, PY [8] |
| 19 | *MED11* | MY, PY | MY, FY, PY [8] |
| 19 | *RAB5C* | MY | PP [1] |
| 19 | *TM4SF5* | MY, PY | MY, FY, PY [8] |
| 19 | *ZMYND15* | MY, PY | MY, FY, PY [8] |
| 19 | *PLD2* | MY, PY | MY, FY, PY [8] |
| 19 | *PSMB6* | MY, PY | MY, FY, PY [8] |
| 19 | *HCRT* | MY | PP [1] |
| 19 | *KCNH4* | MY | LP, PP, LC [24, 40] |
| 19 | *STAT5A* | MY | PP, MY [16, 68] |

| | | | |
|---|---|---|---|
| 19 | *ZNF385C* | MY | LC [40] |
| 19 | bta-mir-195 | PY | MY, FY, PY [8] |
| 19 | bta-mir-497 | PY | MY, FY, PY [8] |
| 19 | *BCL6B* | PY | MY, FY, PY [8] |
| 19 | *RNASEK* | PY | MY, FY, PY [8] |
| 19 | C19H17orf49 | PY | MY, FY, PY [8] |
| 19 | *MINK1* | PY | MY, FY, PY [8] |
| 19 | *ZFP3* | PY | FY, PY [8] |
| 19 | *DERL2* | PY | FY, PY [8] |
| 19 | *DHX33* | PY | FY, PY [8] |
| 19 | *SPAG7* | PY | FY, PY [8] |
| 19 | *ENO3* | PY | FY, PY [8] |
| 19 | *ARRB2* | PY | MY, FY, PY [8] |
| 19 | *ALOX15* | PY | MY, FY, PY [8] |
| 19 | *CAMTA2* | PY | FY, PY [8] |
| 19 | *NLRP1* | PY | FY, PY [8] |
| 19 | *ALOX12* | PY | MY, FY, PY [8] |
| 19 | *ENSBTAG00000013906* | PY | FY, PY [8] |
| 20 | *GHR* | MY, FY | MY, FY, PY, FC, PC, CC, CY, LC, LY [8, 25, 69, 70] |
| 20 | *OXCT1* | MY | MY, FP, PP [20] |
| 20 | *FBXO4* | MY, FY | MY, PP [16, 20] |
| 20 | *ZNF131* | MY | MY [16] |
| 20 | *ENSBTAG00000054352* | MY | MY [16] |
| 20 | *ENSBTAG00000052195* | MY | MY [16] |
| 20 | *RANBP17* | FY, PY | FP [62] |
| 20 | *ENSBTAG00000054687* | PY | MY, FY, PY [71] |
| 20 | *SLC9A3* | PY | MY, FY, PY [71] |
| 20, 27 | 5S rRNA | PY, MY | FY, PY, FP, LP [16, 72] |
| 20 | *AHRR* | PY | MY, FY, PY [71] |
| 20 | *EXOC3* | PY | MY, FY, PY [71] |
| 20 | *NKD2* | PY | MY, FY, PY [71] |
| 20 | *SLC12A7* | PY | MY, FY, PY [71] |
| 20 | *CCDC127* | PY | MY, FY, PY [71] |
| 20 | *SDHA* | PY | MY, FY, PY [71] |
| 20 | *PDCD6* | PY | MY, FY, PY [71] |
| 20 | *PTGER4* | PY | MY [20] |
| 20 | U2 | PY | FY (BTA12), PC (BTA29) [16, 18] |
| 20 | *IRX1* | PY | PY, PP [8, 58] |
| 20 | *PIK3R1* | PY | MY, FY, PY, FP, PP [73] |
| 24 | *PIGN* | MY, PY | MY, PY [74] |

| | | | |
|---|---|---|---|
| 24 | *HRH4* | MY | differentially expressed in blood transcriptome between high and low yielding cows [75] |
| 25 | *SETD1A* | MY | milk synthesis [76] |
| 25 | U6 | MY | milk production [77] |
| 25 | *ALDOA* | MY | milk production [78] |
| 25 | *ITGAD* | FY, PY | MY, FY, PY, FP [78] |
| 26 | *PAX2* | FY | fatty acid [59] |
| 26 | *SNORA70* | FY | fatty acid [59] |
| 26 | *SCD* | FY | fatty acid [59, 79] |
| 26 | bta-mir-12016 | FY | FY [16] |
| 26 | *SLF2* | FY | fatty acid [59] |
| 26 | *HIF1AN* | FY | FY, fatty acid [16, 59] |
| 26 | *LIPK* | PY | fatty acids [80] |
| 26 | *TACC2* | PY | PY, PP [37] |
| 26 | *RNLS* | PY | fatty acids [81] |
| 26 | *DHX32* | PY | FY, PY , FP, PP [74] |
| 26 | *KIF20B* | PY | fatty acids [81] |
| 26 | *PTEN* | PY | mammary gland development and lactation [82] |
| 27 | *GOLGA7* | MY, PY | FY, FP [37] |
| 27 | *GINS4* | MY, PY | FP [16] |
| 27 | *GPAT4* | MY, PY | FC, LP, FP [18, 24] |
| 27 | *ENSBTAG00000027629* | MY, PY | FY, PY [83] |
| 27 | *SFRP1* | MY | FP [84] |
| 27 | *THRB* | FY | LP [49] |
| 28 | *ZMIZ1* | MY, FY, PY | MY, FY, LP [8, 16] |
| 28 | *ZNF365* | MY | MY [16] |
| 29 | *IGF2* | MY, FY | FY, PY, FP, PP [85] |
| 29 | *ASCL2* | MY, FY | FY [16] |
| 29 | *KIRREL3* | FY | MY [86] |
| 29 | *CCDC15* | FY, PY | fatty acid [87] |
| 29 | *NAP1L4* | FY | MY [88] |
| 29 | *PKNOX2* | PY | FY, FP [37] |

**Figure S1 Concordance between imputed KuhVision AF and 1000 Bulls Run9 AF from BTA16**

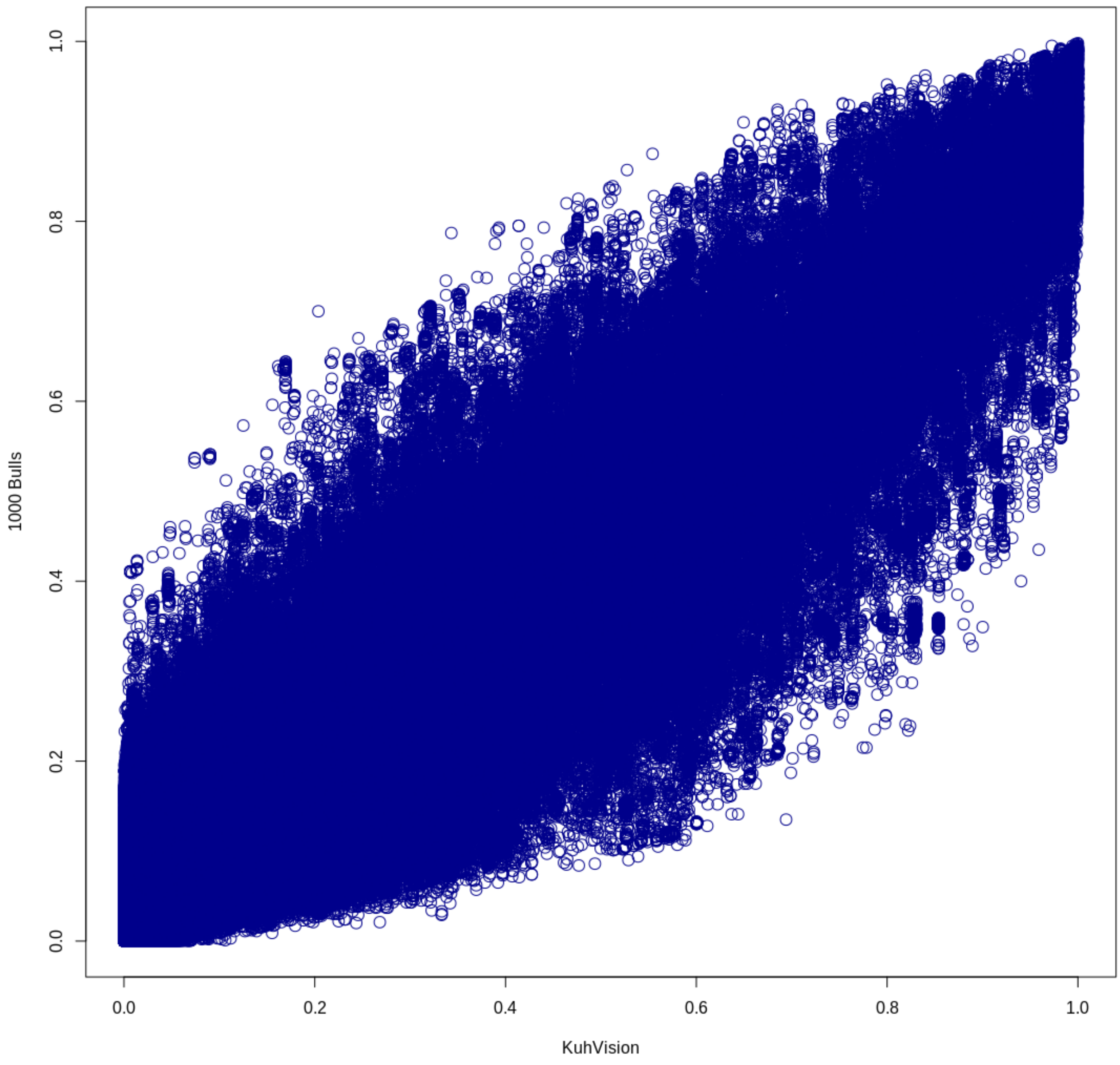

### List of abbreviations

CC – casein content

CY – casein yield

FC – fat content

FP – fat percentage

IGF-1 – insulin-like growth factor-1

LP – lactation persistency

LPC – lactose percentage

LY – lactose yield

PC – protein content

PP – protein percentage

PUFA – polyunsaturated fatty acids

UFA – unsaturated fatty acids

16. Pedrosa VB, Schenkel FS, Chen S-Y, Oliveira HR, Casey TM, Melka MG, Brito LF. Genomewide Association Analyses of Lactation Persistency and Milk Production Traits in

Holstein Cattle Based on Imputed Whole-Genome Sequence Data. Genes (Basel) 2021. doi:10.3390/genes12111830.

38. Li X, Li G, Du X, Sun X, Peng Z, Zhao C, et al. Increased autophagy mediates the adaptive mechanism of the mammary gland in dairy cows with hyperketonemia. J Dairy Sci. 2020;103:2545–55. doi:10.3168/jds.2019-16910.
39. Carrara ER, Dauria BD, Gervásio IC, Silveira RMF, Rovadoski GA, Petrini J, et al. Comparison of Marker Effects and Breeding Values at Two Levels at THI for Milk Yield and Quality Traits in Brazilian Holstein Cows. Genes (Basel) 2022. doi:10.3390/genes14010017.
40. Lopdell TJ, Tiplady K, Struchalin M, Johnson TJJ, Keehan M, Sherlock R, et al. DNA and RNA-sequence based GWAS highlights membrane-transport genes as key modulators of milk lactose content. BMC Genomics. 2017;18:968. doi:10.1186/s12864-017-4320-3.
41. Lu X, Arbab AAI, Abdalla IM, Liu D, Zhang Z, Xu T, et al. Genetic Parameter Estimation and Genome-Wide Association Study-Based Loci Identification of Milk-Related Traits in Chinese Holstein. Front Genet. 2021;12:799664. doi:10.3389/fgene.2021.799664.
42. Song X, Zhao M, Cao Q, Wang S, Li R, Zhang X, et al. Transcriptome provides insights into bovine mammary regulatory mechanisms during the lactation cycle. Journal of Applied Animal Research. 2022;50:275–88. doi:10.1080/09712119.2022.2064865.
43. Friedrich J. Behaviour of cattle – molecular biological background and relationship to milk performance [Dissertation]. Rostock: University of Rostock; 2015.
44. Singh A, Kumar A, Gondro C, Pandey AK, Dutt T, Mishra BP. Genome Wide Scan to Identify Potential Genomic Regions Associated With Milk Protein and Minerals in Vrindavani Cattle. Front Vet Sci. 2022;9:760364. doi:10.3389/fvets.2022.760364.
45. Poulsen NA, Robinson RC, Barile D, Larsen LB, Buitenhuis B. A genome-wide association study reveals specific transferases as candidate loci for bovine milk oligosaccharides synthesis. BMC Genomics. 2019;20:404. doi:10.1186/s12864-019-5786-y.

59. Palombo V, Milanesi M, Sgorlon S, Capomaccio S, Mele M, Nicolazzi E, et al. Genome-wide association study of milk fatty acid composition in Italian Simmental and Italian Holstein cows

using single nucleotide polymorphism arrays. J Dairy Sci. 2018;101:11004–19. doi:10.3168/jds.2018-14413.
